# Variable *Shh* and *Fgf8* positioning in regenerating axolotl limb guarantees consistent limb morphogenesis in different limb sizes

**DOI:** 10.1101/2022.01.04.475010

**Authors:** Saya Furukawa, Sakiya Yamamoto, Rena Kashimoto, Yoshihiro Morishita, Akira Satoh

## Abstract

Limb regeneration in *Ambystoma mexicanum* occurs in various size fields and can recreate consistent limb morphology. The mechanism that supports such stable limb morphogenesis regardless of size is unknown. Sonic hedgehog (SHH) and fibroblasts growth factor 8 (FGF8) play important roles in anteroposterior limb patterning, similar to other tetrapods. Focusing on these two factors, we investigated the detailed expression pattern and function of *Shh* and *Fgf8* in various blastema sizes in axolotl limb regeneration. We measured and functionally analyzed the expression domains of *Shh* and *Fgf8* in regenerating limb blastema of various sizes, and found that, although the position and size of the *Shh*^+^ and *Fgf8*^+^ domains varied depending on the size of the blastema, the secretion of SHH was maintained at a relatively fixed working distance, regardless of blastema size. This stable secretory distance of SHH resulted in the formation of an active proliferative zone (aPZ) in the vicinity of SHH, regardless of blastema size. The aPZ was under the mitogenic influence of SHH and FGF8, resulting in high cell density in the aPZ. We also examined the impact of the aPZ on digit formation. We found that the first digit formation occurs in the aPZ. Next, the aPZ gradually shifts posteriorly as digits develop, which contributes to new digit formation at the site of the shifted aPZ. We also found that the exogenously formed aPZ caused extra digit formation even after the completion of autopod morphogenesis. Our findings suggest that the variable *Shh*-*Fgf8* positioning in various blastema sizes causes various positioning of the aPZ, and that the aPZ leads to digit formation. The mechanism we propose here accounts for stable digit morphogenesis regardless of blastema sizes and urodele-specific digit formation.

**One-Sentence Summary:** A unique SHH-FGF8 spatial interaction compensates for robust limb morphogenesis in various limb sizes.

## Introduction

Urodele amphibians have the ability to regenerate at the organ level. In contrast, amniotes, including humans, have poor regeneration ability. Numerous studies have sought to clarify the mechanisms for the differences in the regeneration ability between urodele amphibians and amniotes. Among them, limb regeneration has been the most selected to study the superior regeneration ability in urodele amphibians.

Limb regeneration accompanies the formation of a regeneration-specific structure called a regeneration blastema. A blastema consists of many undifferentiated cells called blastema cells, some of which are multipotent and some monopotent ^1, 2^. Recent and classic studies indicate that the heterogeneous blastema cell population exhibits similar features and gene profiles to cells in developing limb buds^3, 4, 5, 6^.

A patterned limb is re-created from a blastema and the patterning mechanisms have been considered to be mimicking limb developmental processes. Axolotl limb development has been investigated at the molecular level and the insights become sufficient to compare its process with other tetrapods. Fundamentally, most limb developmental systems are comparable. For example, *Shh* is expressed in the posterior margin of a developing limb bud^7, 8, 9^. The secreted SHH from the margin of the posterior side of a limb bud is received by PTC, which is a SHH receptor and exhibits a relatively wider range of expression domains compared to *Shh*^8^. The posterior-shifted SHH activity regulates the anteroposterior patterning. This anteroposterior patterning controlled by SHH is believed to be conserved among tetrapods. However, an axolotl-specific limb patterning mechanism has also been revealed. *Fgf8* expression is significantly different from that of amniotes ^7, 10^. In amniotes, such as chicks and mice, *Fgf8* expression is restricted to the limb ectoderm during limb development^11, 12^. The ectodermal region where *Fgf8* is expressed is called the apical ectodermal ridge (AER), and it plays a central role in the proximodistal elongation and patterning of a developing limb bud. However, the *Fgf8* expression domain is observable in the mesenchyme in axolotl limb buds ^14^. How this axolotl-specific gene expression pattern contributes to axolotl limb development remains unknown. In addition to the gene expression pattern, limb patterning also exhibits axolotl specificity. In amniotes, the limb digits are basically formed in a posterior to anterior order ^13, 14^. In contrast, axolotls form digits in the reverse order ^15^. This digit patterning mechanism has remained unclarified and has been a missing link in the evolutionary study. In limb regeneration, these conserved and specific features in limb development are considered to be faithfully recreated in a regenerating blastema. To date, similar gene expression patterns have been reported between a developing limb and a regenerating blastema ^4^. Furthermore, it was revealed that a formed axolotl blastema could contribute to limb development when a part of the blastema was grafted into a developing limb bud ^5^. This indicates the similar properties between a developing limb bud and a regenerating blastema. In this context, blastema induction can be considered to be a rewinding of the developmental clock.

Although limb regeneration has been considered a recapitulation of limb developmental processes in most processes, there are certainly major differences between them. One of the differences between them should be the size. Limb amputation is performed with limbs that are much larger compared to a developing limb bud. Subsequently, a blastema, which has been considered as a similar structure to a developing limb bud, is formed on the amputation plane. Then, the induced blastema undergoes limb morphogenesis. Thus, the field of morphogenesis in limb regeneration is much larger than that in limb development. According to the variation in field size, gene expression should be adapted to the size. However, gene expression in varied field sizes has not yet been investigated in detail.

In the present study, we focused on the axolotl-specific gene expression pattern in varied blastema sizes. Our findings suggest that the gene expression patterns specific to urodele amphibians play an important role in supporting the successful limb regeneration that occurred in a variety of sizes. We also demonstrate that the axolotl-specific regulation of the *Shh* and *Fgf8* expression in a regenerating blastema leads to axolotl-specific digit development with reversed order. Our results provide a new perspective on the consistent regenerative capacity of urodele amphibians in various sizes of a field and insights into the missing link in the evolutionary study of digit morphogenesis.

## Results

### Measurement of the regeneration field in axolotl limbs

Axolotls can regenerate their limbs regardless of their size. In laboratory conditions, limb regeneration is usually observable in a variety of animal sizes (Fig. 1A–C). Larger animals take a longer time to complete limb regeneration, and smaller animals take a shorter time (Fig. 1A–C). This variation in timing and size makes a determination of the blastema stage difficult. In this study, we determined the blastema stage based on blastema appearance, characterizing a late blastema by dorsoventral flattening and a ratio of anterior-posterior to the proximal-distal length of approximately 1:1. The outlines of each limb and blastema are shown in Fig. 1D to compare the size. Though there are differences in regeneration time, axolotls can regenerate their limbs throughout their life.

**Fig. 1.**
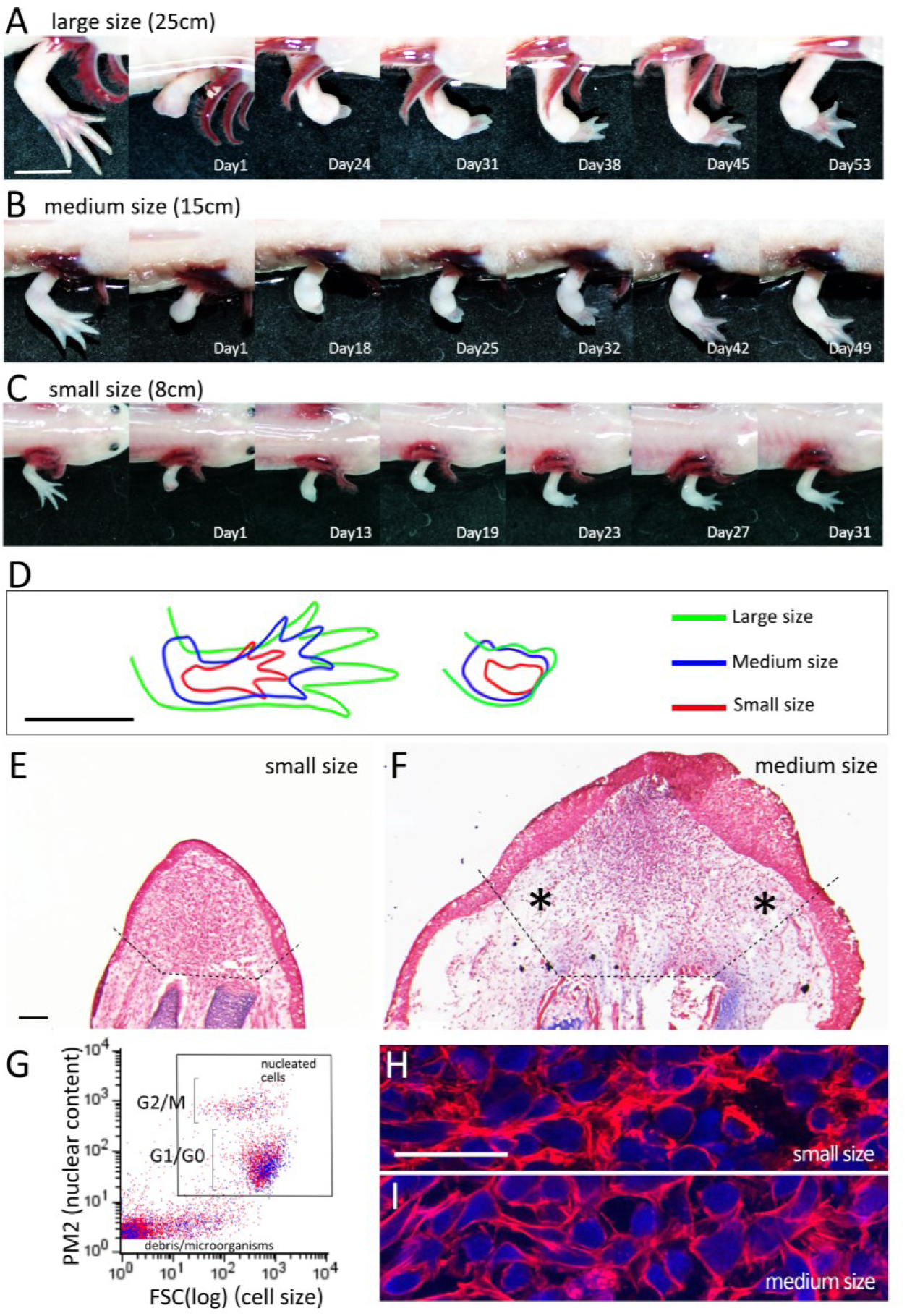
Limb regeneration at different animal sizes and blastema formation. (A–C) Time course of the limb regeneration process in different size animals. Scale bar in A = 1 cm. (D) The outlines of the limbs and regeneration blastemas are shown in A–C. The regeneration blastemas were chosen to have a 1:1 anteroposterior width and proximodistal width at each size. Scale bar in D = 1 cm. (E, F) Alcian, eosin & hematoxylin staining. The dotted lines indicate the presumptive amputation plane. Asterisks indicate the sparse regions. Histological analysis reveals the presence of the sparse region in the larger size of the blastema. Scale bar in E = 200 µm. The Posterior is to the left. (G) Flow cytometric analysis. The red and the blue dots indicate the blastema cells derived from medium- and small-size animals, respectively. No significant difference in cell size was observed. (H) Phalloidin rhodamine and Hoechst staining. Scale bar in H = 50 µm.

We histologically compared blastemas of different sizes and investigated the cell size. For experimental convenience, we selected small- and medium-sized animals for most experiments. We obtained similar histological results in both sections but identified a few differences. The blastemas that grew in medium-size animals (BL-M) exhibited the low-cell density in the anterior and posterior regions (Fig. 1F, asterisks). The blastema epithelium of BL-M tended to be thicker than that in the blastema of small-size animals (BL-S) (Fig. 1E, F). We also compared the cell size in BL-S and BL-M using flow cytometry (Fig. 1G). Blastemas were enzymatically digested and then underwent flow cytometric analysis directly. The signal of the forward scatter (FSC) of the flow cytometry results indicates cell size. Within the nucleated population (PM2^+^), the plots from BL-M and BL-S were distributed in the same areas. The similarity of the cell size in BL-M and BL-S was also confirmed on the sections, in which the outlines of the cells were visualized by phalloidin rhodamine (Fig. 1H, I). These suggest that the cell size was consistent throughout their life although the size difference cause histological differences between BL-S and –M.

### Shh and Fgf8 expression domains in different blastema sizes

We next investigated how genes were expressed in blastemas of different sizes. *Shh* and *Fgf8* expression patterns were investigated by *in situ* hybridization since *Shh* and *Fgf8* are known to play an essential role in limb patterning in many animals (Fig. 2). In BL-S, *Shh* was typically expressed locally at the margin of the posterior mesenchyme along with the blastema epidermis (Fig. 2A, B). In BL-M, *Shh* expression appears to be regulated differently at the proximodistal level. In the distal region of BL-M, the *Shh^+^* domain was restricted in the posterior margin (Fig. 2F). The *Shh^+^* domain of the proximal side, however, tends to be apart from the posterior margin of the blastema epithelium (Fig. 2E, F). It is of note that the disconnection of the *Shh^+^* domain from the posterior margin is a characteristic expression pattern. *Fgf8* was expressed in the anterior side of the blastema mesenchyme as described ^13^(Fig. 2C, D, G, H). As shown in Fig. 2C and G, *Fgf8* was expressed in the anterodorsal and anteroventral blastema mesenchyme. The longitudinal sections were prepared with a slight deviation from the midline due to this characteristic *Fgf8^+^* domain (Fig. 2B, F, D, H). The detachment of the *Fgf8^+^* domain from the anterior margin was commonly observed (Fig. 2C, G). To describe the *Shh^+^* and *Fgf8^+^* domain in more detail, a three-dimensional (3D) reconstitution was performed (Fig. 3, Sup. Mov. 1 and 2). Two sets of serial sections, one each for *Shh* and *Fgf8*, underwent regular *in situ* hybridization procedures. Then, all sections were captured and captured all images were reconstituted into a 3D image (Fig. 3). In the BL-S, the *Shh*^+^ domain was at the posterior margin of the mesenchyme (Fig. 3C, E, F, H). The *Fgf8*^+^ cells could be detected from the relatively distal portion of the blastema mesenchyme (Fig. 3H, N). The *Fgf8*^+^ domain and the *Shh*^+^ domain were close by but appear not to be merged at any level (Fig. 3H). In the BL-M, the position of the *Shh*^+^ domain was relatively variable as mentioned below. Figure 3 shows one representative example of the *Shh*^+^ and *Fgf8*^+^ domains in the BL-M. The *Shh*^+^ and *Fgf8*^+^ domains were observable in the posterior and anterior regions of the blastema, respectively (Fig.3I-N). *Fgf8*^+^ cells were located relatively middle region of the blastema as compared to the BL-S (Fig. 3G and M). *Shh*^+^ domain is located along the posterior margin in the distal region, but apart from the margin in the posterior region (Fig. 3L). Overall, the *Shh*^+^-*Fgf8*^+^ domain can be found as being posteriorly shifted in most blastema. The spatial expression pattern of *Shh* and *Fgf8* in the BL-S and the BL-M clarifies the similarity and dissimilarity of those in the different sizes of the blastema.

**Fig. 2.**
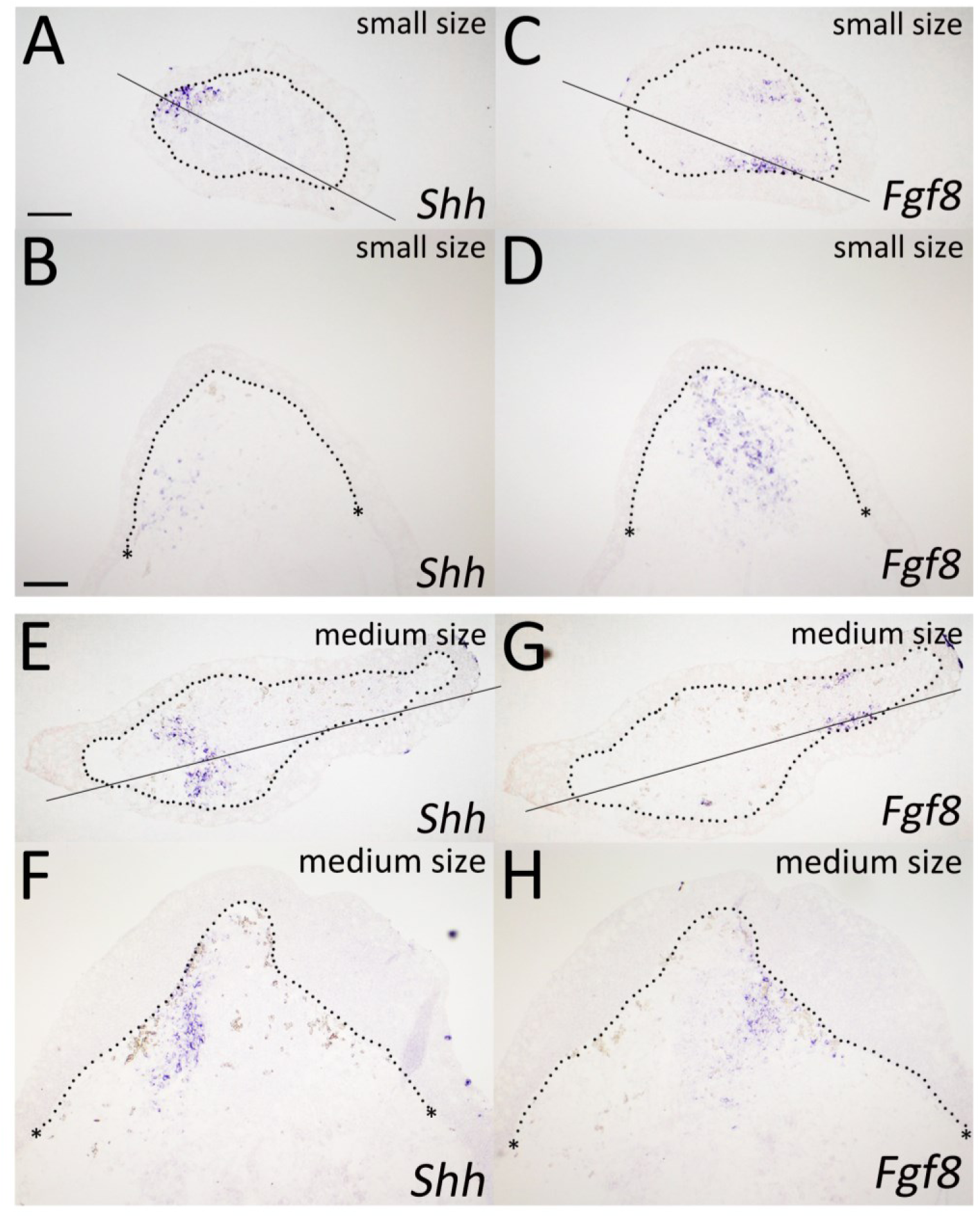
*Shh* and *Fgf8* expression pattern in the BL-S and -M. (A–H) *Shh* and *Fgf8* expression was visualized by *in situ* hybridization in the BL-S (A–D) and BL-M (E–H). The dotted lines indicate the border of the blastema epithelium. Asterisks denote the punctuated dermal collagen, indicating the amputation plane. Lines in A, C, E, and G indicate the approximate position of the longitudinal sections shown in B, D, F, and H, respectively. Scale bars in A and B = 200 µm. The Posterior is to the left (A–H).

**Fig. 3.**
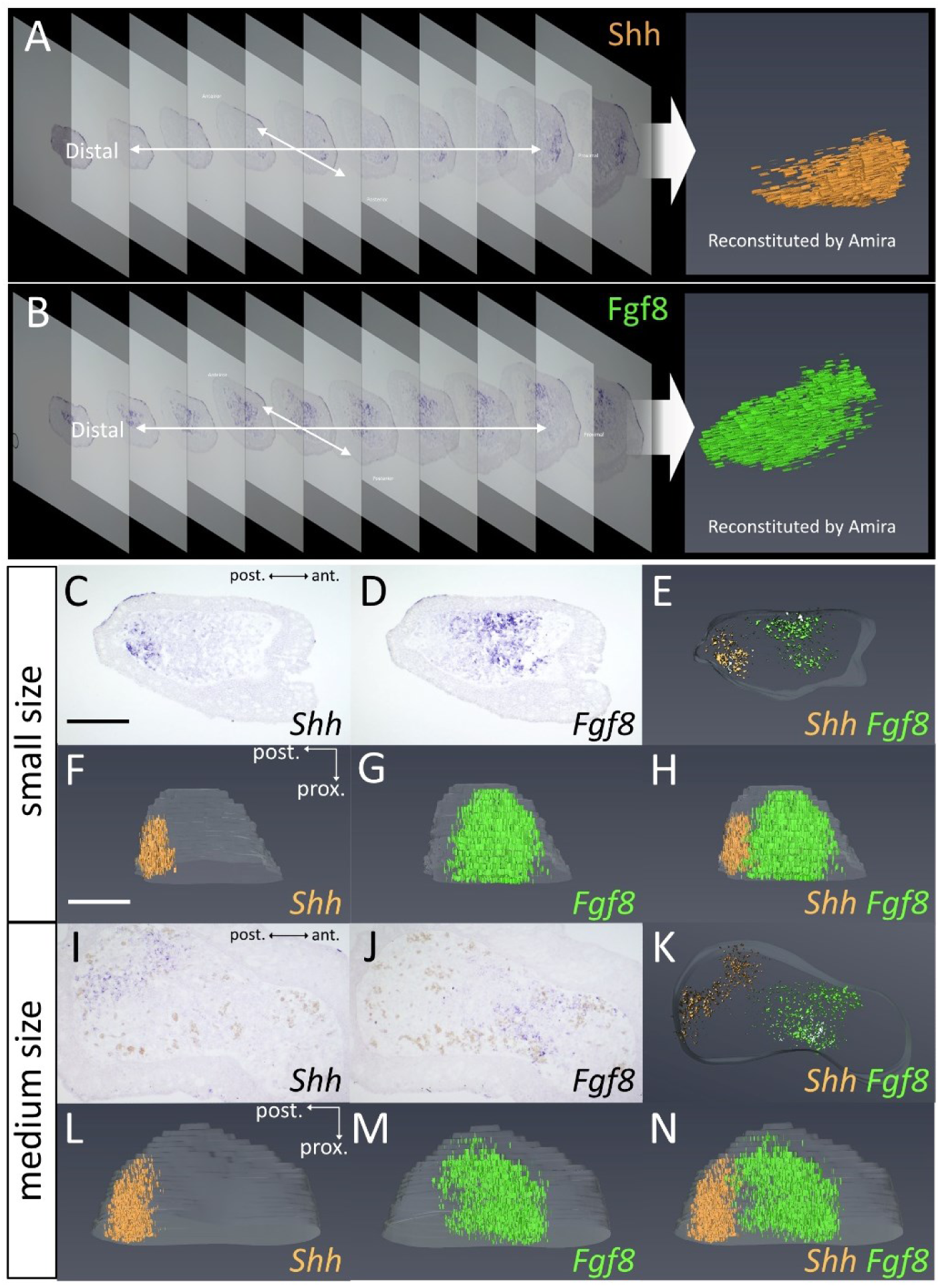
The 3D reconstructed image of the *Shh^+^* and *Fgf8^+^* domain in the BL-S and -M. (A, B) The schematic diagram of the creation of the 3D image of *Shh* and *Fgf8* expression. The signals of *Shh* or *Fgf8* cells were extracted from the sections and reconstructed into a 3D image. (C–H) Gene expression pattern in the BL-S. Typical *Shh* and *Fgf8* expression patterns in the BL-S are shown in C and D, respectively. E–H are the reconstituted images showing *Shh* (orange) and *Fgf8* (light green) expression. (E) The proximal view of the proximal six sections. E, F, G, and H are snapshots from the Supplemental movie 1. (I–N) Gene expression pattern in the BL-M. (K) The proximal view of the proximal six sections. K, L, M, and N are snapshots from the Supplemental movie 2. Scale bars in C and F = 400 µm. The Posterior is to the left (C–N).

To investigate further, we measured the *Shh^+^* and *Fgf8^+^* domains in the transverse sections of the blastema (Fig. 4). The measurement scheme is described in Sup. Fig. 1A and B. As described, the *Shh^+^* domain was varied at the proximodistal level (Fig. 2F). To normalize this variation, we selected two or three sections from an identical blastema, on which *Shh* and *Fgf8* expression could be confirmed, and its average was calculated. The average widths are aligned and presented graphically in Fig. 4A and D. (The raw data is shown in Sup. Fig. 1C and D.) The *Shh^+^* domains were located on the posterior side. As illustrated in Fig. 2 and 3, the *Shh^+^* domain tends to be apart from the posterior end of the blastema mesenchyme in larger blastemas (Fig. 4A). The scatter diagram of the width of the *Shh^+^* domain and the blastema width shows a clear correlation (R = 0.8248) between them (Fig. 4B). Focusing on the occupation ratio of the *Shh^+^* domain in the blastema, the ratio of the *Shh^+^* domain was relatively constant, at an average of 22.8%. (Fig. 4C). No difference in the *Shh^+^* domain between the forelimb and the hind limb could be found (Fig. 4B, C). As for the *Fgf8^+^* domain, the *Fgf8^+^* domain was wider than the *Shh^+^* domain and was located on the anterior side of the blastemas (Fig. 4D). The *Fgf8^+^* domain was much weakly correlated with the blastema as compared to the *Shh*^+^ domain (Fig. 4E). On the other hand, it was observed that the range of *Fgf8* expression relative to the size of the regenerating blastemas tended to be constant regardless of size, although there was a larger variation as compared to the *Shh*^+^ domain (Fig. 4F). Lastly, we aligned the *Shh^+^* and *Fgf8^+^* domains on the adjacent sections (Fig. 4G). On most sections, the *Shh^+^* and *Fgf8^+^* domains were not merged, and a certain distance existed between the two expression domains (Fig. 4G). Those results indicate the expression domain of *Shh* and *Fgf8* are scaled along with the blastema size and the position of *Shh*^+^ and *Fgf8*^+^ domains are varied in the blastema mesenchyme.

**Fig. 4.**
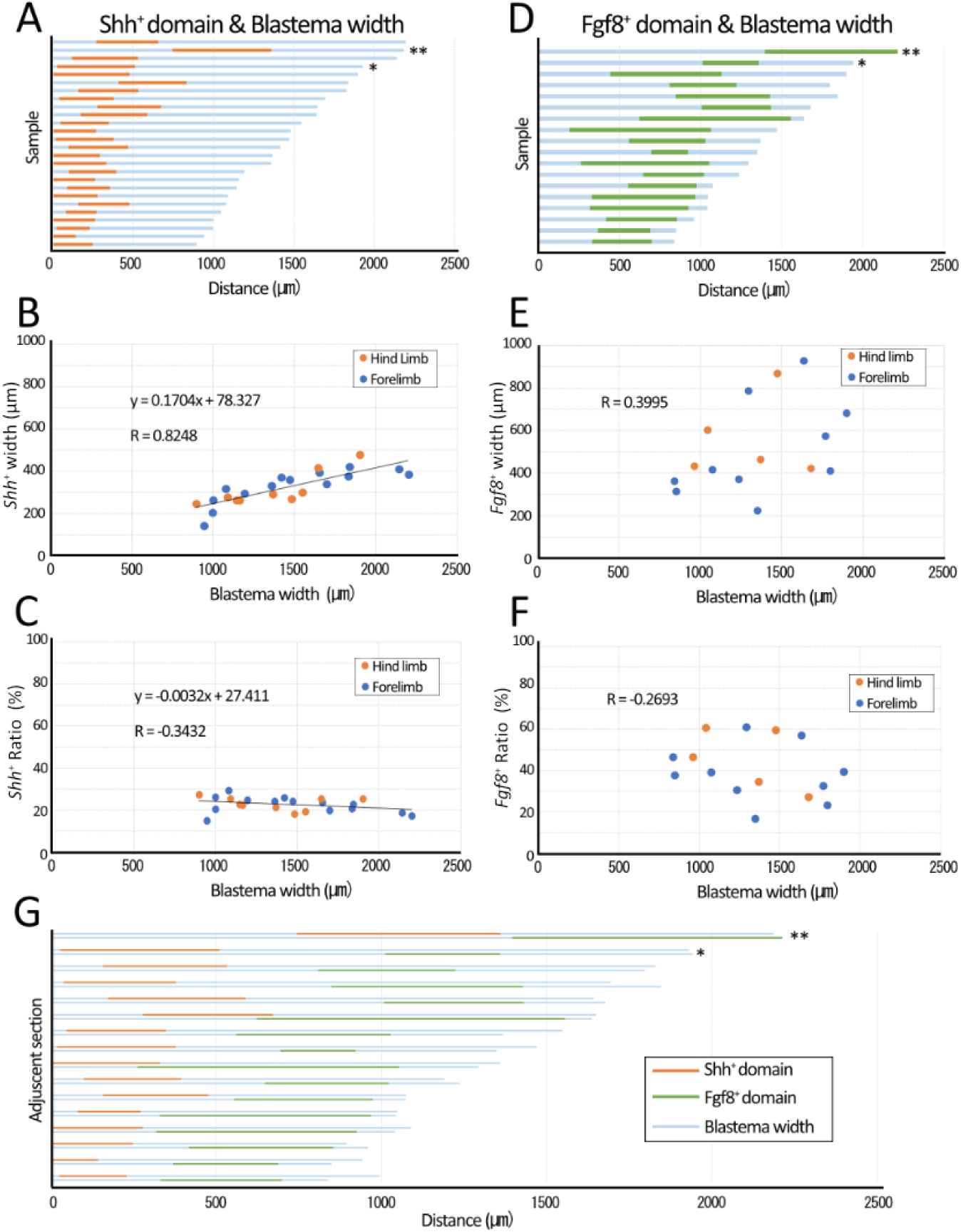
Measurement of the *Shh^+^* and *Fgf8^+^* domain in blastemas. (A, D) The averaged width of the *Shh*^+^ and *Fgf8*^+^ domains collected from two to three sections is shown in orange and green, respectively. Light blue indicates the width of the blastema mesenchyme along the anteroposterior axis. (B, E) The scatter plot illustrating the relationship between the width of the *Shh*^+^ or *Fgf8*^+^ domain and the blastema width. The line in B shows the collinear approximation represented by the formula. R shows the correlation coefficient. (C, F) The scatter plot illustrating the relationship between the ratio of the *Shh*^+^ or *Fgf8*^+^ domain in the blastema mesenchyme and the blastema width. The line in C shows the collinear approximation represented by the formula. R shows the correlation coefficient. (G) Two serial sections were prepared and *in situ* hybridization was performed on each series of sections to visualize the *Shh*^+^ and *Fgf8*^+^ cells. Each expression domain was measured and visualized in the same way as in (A) and (D). Only the samples that could be measured in the identical sample were extracted and shown. The asterisks and the double-asterisks in A, D, and G indicate the samples used for BrdU and cell density analysis in Fig. 7I-P and Fig. 7Q-X, respectively.

We also investigated whether the expression of *Shh* and *Fgf8* is mutually dependent. We kept animals with blastemas at the mid-bud stage in water containing SU5402 (as an Fgf-signaling inhibitor), or cyclopamine (as an Shh-signaling inhibitor). The blastemas were harvested after 3 days of housing with the chemical compound. Then, the *Shh* and *Fgf8* expression levels were measured by quantitative RT-PCR (Sup. Fig. 2A). Both chemical compounds decreased in the expression level of *Shh* and *Fgf8*. A similar result was reported using axolotl limb buds, thereby supporting the results. These findings suggest that *Shh* and *Fgf8* are maintained in a mutually dependent manner.

Next, we investigated how the distribution of secreted factors was affected by gene expression that is scaled along with blastema size, focusing on *Shh*, which plays a major role in the anteroposterior limb patterning. The *Ptc1* gene encodes a receptor for SHH and its expression has been used as an indicator of the SHH distribution^16, 17^. *Shh* is expressed in the posterior margin of an amniote limb bud and SHH is secreted toward the anterior side, creating an SHH gradient in the limb bud ^18, 19^. As in the amniote limb bud, *Ptc1* was found in the posterior side of the BL-S and BL-M (Fig. 5B, D, Sup. Fig. 1F, H). The *Ptc1^+^* domain was overlapped with the *Shh^+^* domain and extended to the anterior, forming a relatively wider expression domain compared to the *Shh^+^* domain (Fig. 5A–D, Sup. Fig. 1E-H). As with the *Shh^+^* domain, the *Ptc1^+^* domain width exhibits a correlation with the blastema width (R = 0.7557, Fig. 5E), and the ratio shows consistency (Fig. 5F, average = 40.3%). No apparent differences in the *Ptc1^+^* domain between the forelimbs and the hind limbs were observed. To estimate SHH diffusion, the extent of *Ptc1* expression was investigated. *Ptc1* expression has been used as an indicator of SHH diffusion in limb developmental studies^16, 17^. The lines were drawn at the anterior limits of the *Ptc1^+^* and *Shh^+^* domains, respectively. Next, the width of these two lines (the anterior borders of the *Shh^+^* and *Ptc1^+^* domains) was calculated and defined as the SHH diffusion range (Fig. 5G, H). The SHH diffusion range was constant, suggesting the same distribution of SHH from the anterior border of the *Shh^+^* domain, regardless of blastema size. This constant SHH-diffusion range was also supported by simulations. We simulated the spatial distributions of SHH in tissues with two different sizes of *Shh^+^* domains using a mathematical model for SHH diffusion and degradation (Fig. 5I–K). It was assumed that all cells have the same expression level and secretion rate of SHH in the expression domain. The curves in the figures show the results for different degradation rates. To compare the SHH diffusion ranges between the cases with different sizes of *Shh^+^* domains, the spatial patterns of SHH were shifted in parallel so that the anterior ends of expression domains coincide (Fig. 5K). It was found that the SHH diffusion range was very similar, independent of its expression domain size, although the difference was slightly clearer when the degradation rate was smaller. These findings strongly suggest that the working range of SHH is constant, regardless of the size of the *Shh*^+^ domain and blastema size.

**Fig. 5.**
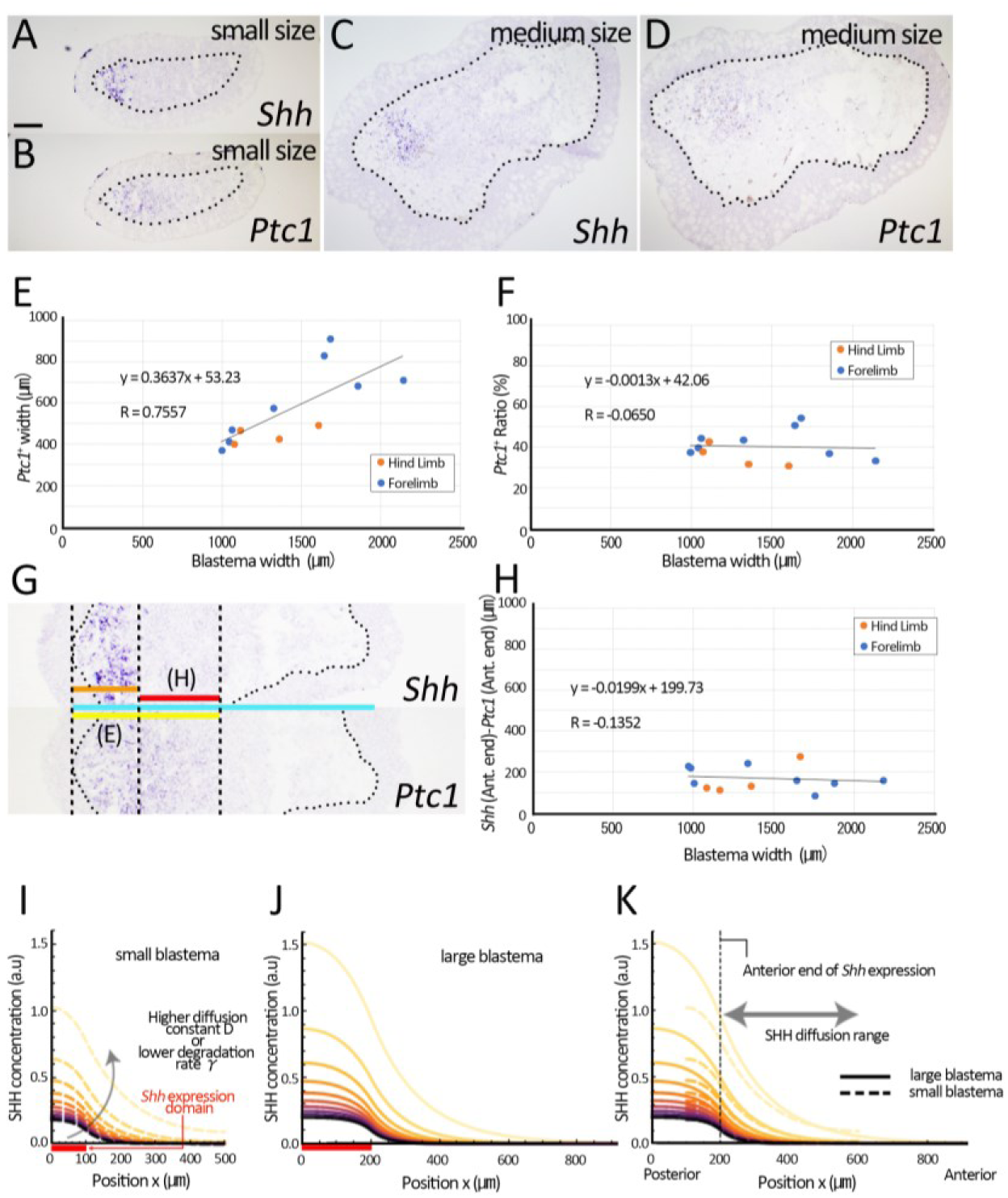
*Ptc1* expression and assumptive SHH distribution in blastemas. (A–D) *Ptc1*or *Shh* expression was visualized by *in situ* hybridization. The dorsal is up and posterior to the left. Scale bar in A = 200 µm. The dotted lines indicate the border of the blastema epithelium. The Posterior is to the left. (E–F) The scatter plot illustrating the relationship between the width or ratio of the *Ptc*^+^ domain and the blastema width. The lines show the collinear approximation represented by the formula. R shows the correlation coefficient. (G) An explanation of the measurement for E and H. The red line indicates the SHH distribution, which was defined as the distance between the anterior borders of the *Shh*^+^ and Ptc1^+^ domain. (H) The scatter plot illustrating the relationship between the assumptive SHH distribution and the blastema width. (I–K) The graphs illustrate the results of simulations of SHH distribution in different widths of the *Shh*^+^ domain. (I) The case of the smaller (100 µm, red line) *Shh*^+^ domain. (J) The case of the larger (200 µm, red line) *Shh*^+^ domain. (K) The merged images aligned with the anterior end of *the Shh* expression. The lines with yellow to purple colors indicate the SHH concentration with different parameters in the diffusion constant (D) and the degradation rate (γ).

### Shh and Fgf8 regulation and functions in regenerating blastemas

We focused on the relationship between the spatial *Shh* expression pattern and the cell density to investigate the function of SHH, which is diffused with a constant range, on blastema cells (Fig. 6). *Shh* expression was visualized by sectional *in situ* hybridization, and the section was subsequently compartmentalized into a 100 µm square by Photoshop CS6 software (Fig. 6A). All cells were counted and all *Shh^+^* cells were marked based on the Hoechst signal. Then, the cell density was visualized (Fig. 6B). The alignment between samples was performed at the anterior end of the *Shh^+^* cell–containing square. The alignment of the data clarified that the higher cell density squares were located around the anterior border of the *Shh^+^* domain. To evaluate this in a three-dimensionally manner, the sections were reconstituted in Amira software (Fig. 6C–F, Sup. Data 1 and 2, Sup. Mov. 3). The nuclei of the blastema mesenchyme were plotted manually based on the Hoechst signal, and the 3D address was obtained. Based on the acquired 3D address, the 3D image was reconstituted as dots. We also marked the *Shh*^+^ cell and displayed it in pink (Fig. 6C, E, Sup. Mov. 3). Cell density was calculated as the average distance from the five nearest nuclei, and shown as the heatmap (Fig. 6D, F, Sup. Data 1 and 2). The density of the cells tends to be relatively higher in the anterior region of the *Shh* expression domain (Fig. 6D, Sup. Data 1). To visualize the cell density on an X-Y plane, we extracted three images from the proximal region and show the proximal view (Fig. 6E, F, Sup. Data 2). The top 10 cells with the highest cell density are roughly concentrated in two areas of the regenerating blastema (Fig. 6F, arrows). One of the two areas, located near the *Shh*^+^ domain, appears to have a higher density compared to the other area. This result suggests that a higher cell density is formed near the *Shh*^+^ domain where SHH is secreted with a constant working distance regardless of blastema size.

**Fig. 6.**
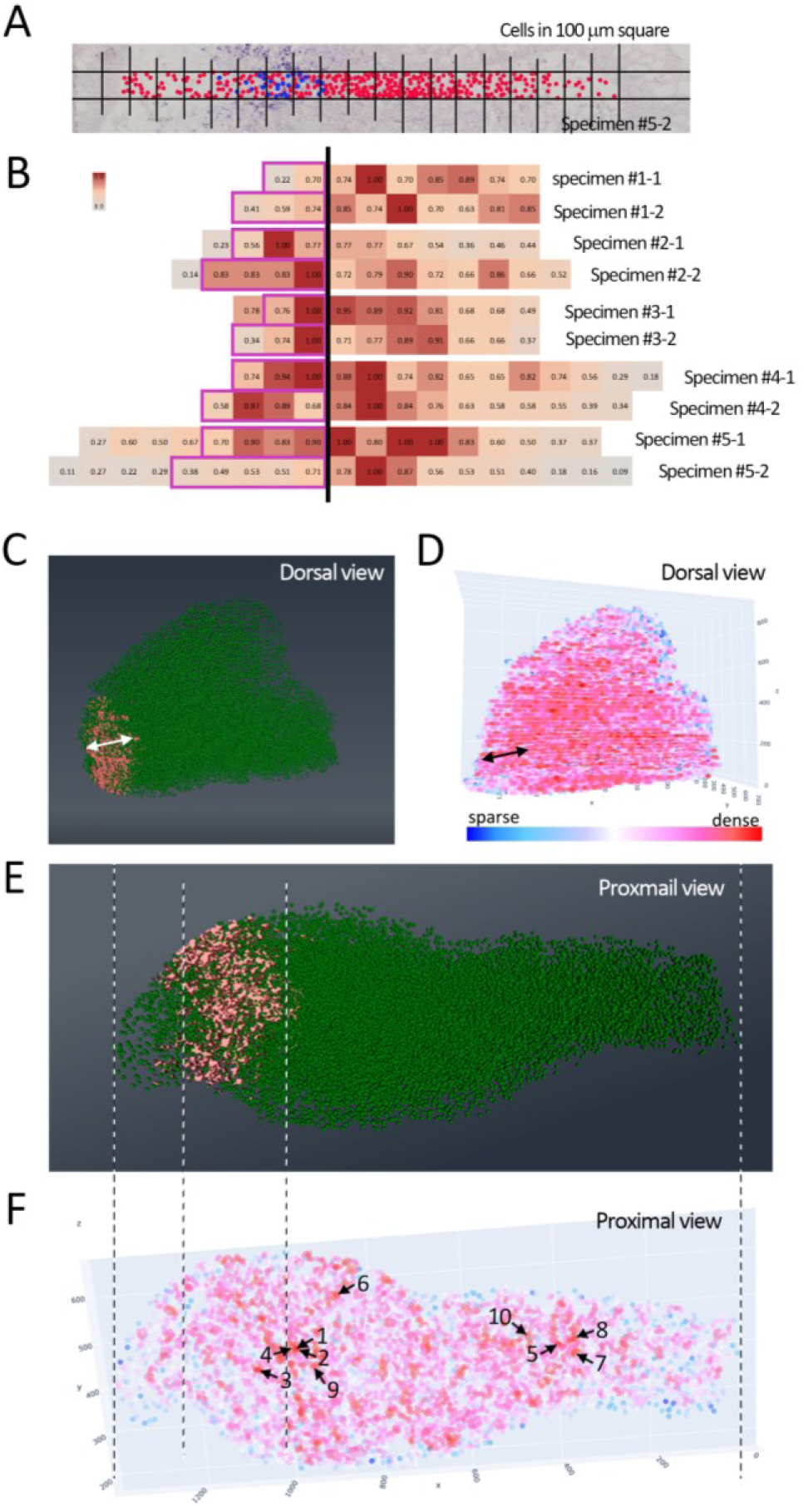
Cell density in regenerating blastema. (A) Cell density and *Shh*^+^ cells in a 100 µm squares were manually counted on the sections. Red and blue dots indicate cells/*Shh*^-^ and cells/*Shh*^+^, respectively. (B) The heatmap graph based on the count as in A. The number of cells in the square with the highest number of cells was set as 1, and the ratio was quantified. The purple boxes show the square containing *Shh*^+^ cells. The bold black line indicates the anterior border of the *Shh*^+^ domain. (C, E) The reconstituted image from the sections. The positions of the nuclei are shown in green, and the *Shh*^+^ signal by *in situ* hybridization is shown in pink. (D, F) The heatmap illustrating cell density in a blastema. The double-headed arrows in C and D indicate the *Shh*^+^ domain in the identical sample. (E, F) Three proximal sections were extracted and observed from the proximal side. Arrows and numbers in E indicate the 10 cells with the highest cell density from the visualized heatmap. The Posterior is to the left (A–F).

Next, we investigated whether the increase in cell density in the vicinity of the *Shh*^+^ domain is relevant to the cell proliferative activity. Two independent series of sections were made from the same sample to visualize gene expression, cell proliferation and cell density, and were three-dimensionally reconstituted with Amira software (Fig. 7). From the reconstituted 3D image, six sections were extracted from the region where *Shh* and *Fgf8* expression could be observed and the image was visualized from the proximal view. Cell proliferation activity was visualized by BrdU incorporation (Fig. 7C, K, S). To visualize the cell density, we made a heatmap using the Distance Map function of Amira software. To focus on the mesenchymal cells, we manually dissected the signal in the epidermis on the software by referring to the Hoechst signal. In the BL-S, *Shh*^+^ and *Fgf8*^+^ domains were observed as described previously (Figs. 2 and 3). Cell proliferation could be observed in all mesenchymal regions (Fig. 7C, E). As evident from the heatmap, higher cell density can be observed in the center region of the mesenchyme, including the area between the *Shh*^+^ and the *Fgf8*^+^ domain (Fig. 7F-H). Regarding the BL-M, the *Shh*^+^ and *Fgf8*^+^ domain is relatively variable as shown in Fig. 4. We selected two representative blastemas; one has the *Shh*^+^ domain at the relatively posterior end of the mesenchyme (Fig. 7I-P) and the other has the *Shh*^+^ domain near the middle portion of the blastema mesenchyme (Fig. 7Q-X). Accordingly, the *Fgf8*^+^ domain was also moved (Fig. 7J, R). The synchronized pattern of BrdU incorporation could be observed (Fig. 7K, S). As the *Shh*^+^-*Fgf8*^+^ domain is located posteriorly (Fig. 7I-P), the higher proliferative activity could be found posteriorly in the blastema (Fig. 7K, M). In the anterior side of the mesenchyme, a smaller number of the BrdU-incorporated cells could be found (Fig. 7K, M). Higher cell density could be observed between the *Shh*^+^ and *Fgf8*^+^ domain (Fig. 7N-P). As shown in Fig. 4, some blastemas have an anteriorly-shifted *Shh*^+^ and *Fgf8*^+^ domain. As the *Shh*^+^-*Fgf*8^+^ domain is located anteriorly (Fig. 7Q-X), the higher proliferative activity could be found anteriorly (Fig. 7S, U). The heatmap also shows that the area with higher cell density was anteriorly shifted, and was located between anteriorly shifted the *Shh*^+^-*Fgf8*^+^ domain (Fig. 7V-X). In this blastema, a sparse area is visible on the posterior side (Fig. 7S, T, U). From these observations, we defined the area that shows higher cell proliferation activity and cell density in axolotl limb blastema as the active proliferation zone (aPZ). The data suggest that the aPZ is formed in the area flanked by *Shh*^+^ and *Fgf8*^+^ domains and that the aPZ is variably formed within a blastema, depending on the variable location of *Shh-Fgf8* expression.

**Fig. 7.**
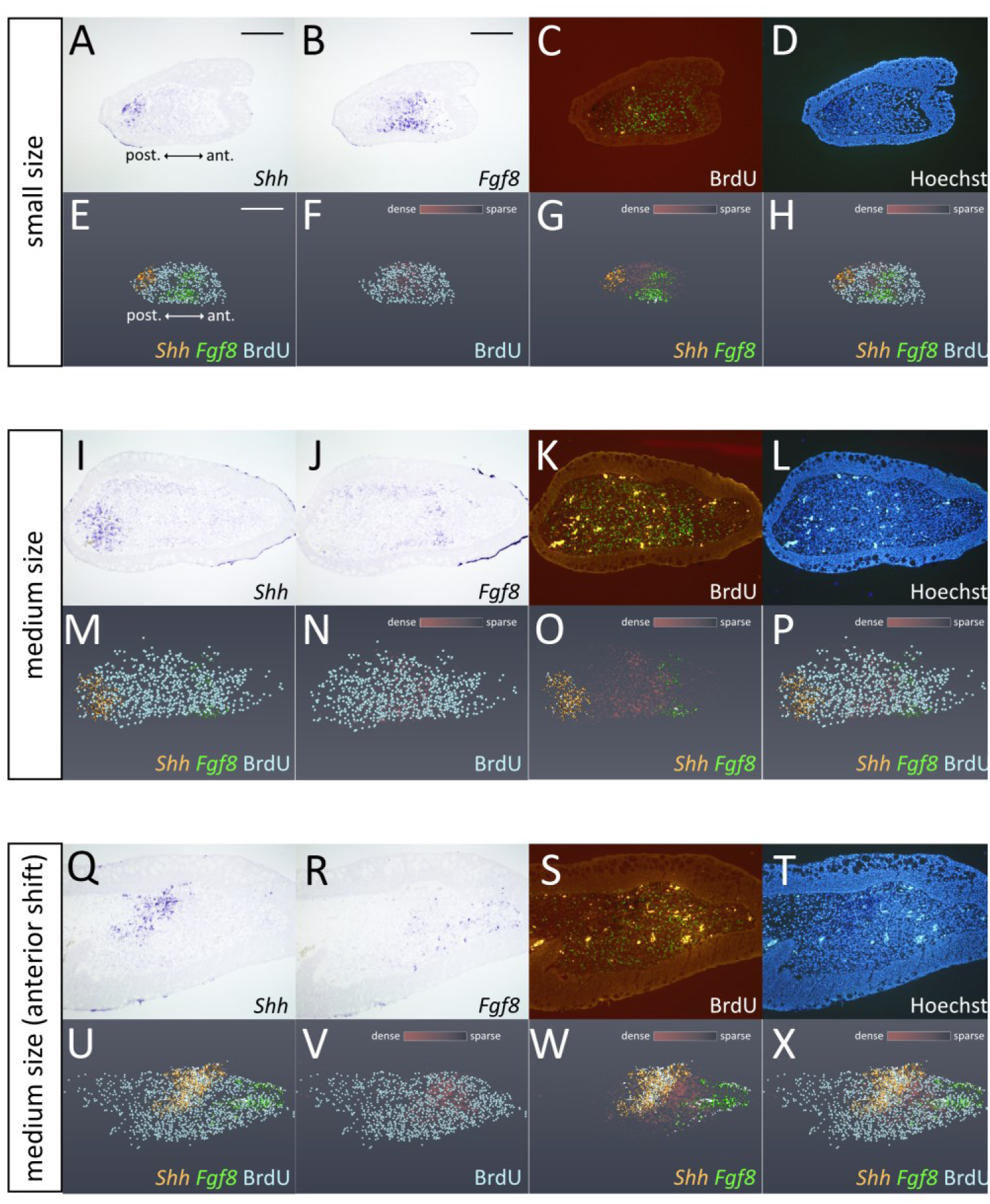
Cell proliferation activity and cell density in the BL-S and BL-M. (A, B, I, J, Q, R) *Shh* or *Fgf8* expression was visualized by *in situ* hybridization. (C, D, K, L, S, T) BrdU incorporation was visualized by immunofluorescence, and nuclei were visualized by Hoechst. A, C, and D are identical sections and B is the adjacent section. Similarly, I, K, and L are identical sections, as are Q, S, and T. (E–H, M–P, U–X) The signals of those in the blastemas were acquired and reconstituted into the 3D image (Sup. Mov. 4–6). The proximal view of the six sections including the section shown in A–D, I–L or Q–T. Each color indicates the following: *Shh* = orange, *Fgf8 =* light green, BrdU = light blue. The cell density is shown as the heatmap (higher cell density = deep brown, lower cell density = deep gray). (A–H) The BL-S. (I–P) The typical BL-M. (Q–X) The BL-M with the anteriorly-shifted aPZ. Scale bar in A and B = 400 µm, in E = approx. 400 µm

We also investigated the mitogenic activity of SHH and FGF8 for blastema cells. We cultured axolotl blastema mesenchymal cells and treated them with SHH and/or FGF8 for 3 days. The *proliferating cell nuclear antigen* (*Pcna*) expression level was measured by quantitative RT-PCR (Sup. Fig. 2B). Solo-application of SHH or FGF8 onto the cultured axolotl blastema cells increased the PCNA expression level compared to the control cells. This is considered to be a similar result as that of the previously published paper using axolotl developing limb bud cells ^8, 20^. We further investigated the cooperative mitogenic activity of SHH and FGF8 onto the cultured blastema cells. Applying both SHH and FGF8 onto the cultured blastema cells simultaneously increased the level of PCNA expression more than that in other conditions. This suggests that the area between the *Shh*^+^ and *Fgf8*^+^ domains has a highly proliferative environment since it can be expected to be exposed by both SHH and FGF8.

Next, we performed a functional analysis to investigate whether SHH has an impact on aPZ formation in a blastema. We attempted to change the *Shh*^+^ domain by electroporation. First, we confirmed that the electroporation itself did not change the pattern of gene expression and cell proliferation by electroporating the control vector, pCS2-AcGFP (Fig. 8A-J). The electroporation of the control vector did not change the *Shh* or *Fgf8* expression pattern in the blastema and did not have a severe impact on the pattern of BrdU-incorporated cells (Fig. 8A-J). As the vector, pCS2-Shh-p2a-AcGFP was electroporated into the blastema, and the new *Shh*^+^ domain emerged in the electroporated region (Fig. 8K, P, Q). The newly formed *Shh*^+^ domain was in the relatively anterior side of the blastema (Fig. 8K, P, Q). The *Fgf8*^+^ domain appeared to have shifted anteriorly and another aPZ was formed. Higher mitotic activity and cell density could be found between the newly formed *Shh*^+^ and *Fgf8*^+^ domain (Fig. 8S, T). The area, which is beside the original *Shh*^+^ domain, becomes less proliferative. These findings suggest that the position of the aPZ is variable in the blastema, according to the positioning of the *Shh*^+^-*Fgf8*^+^ domain.

**Fig. 8.**
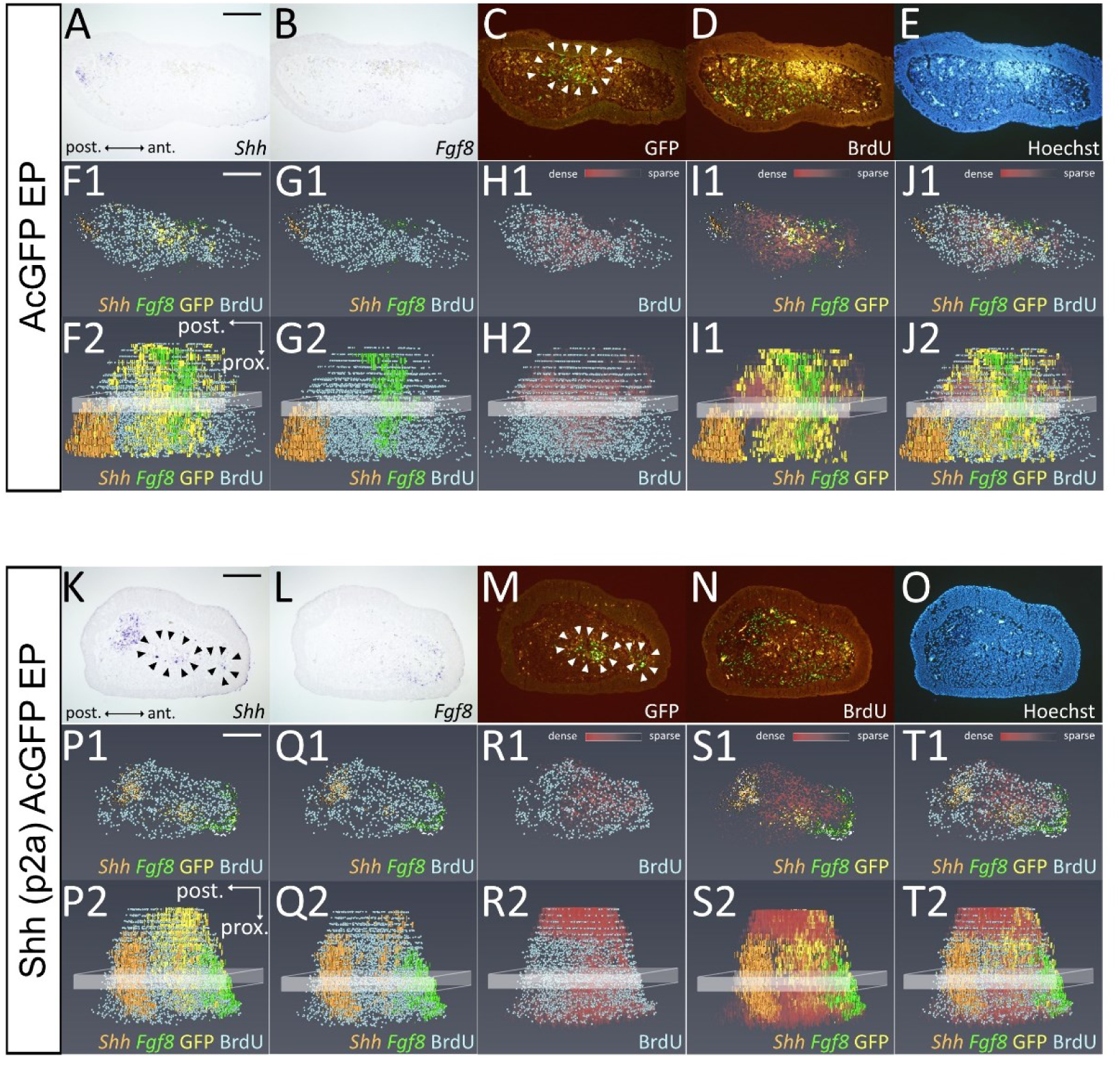
The variability of the positioning of the aPZ caused by ectopic *Shh* expression. Three series of sections were made from the identical blastema. *Shh* and *Fgf8* expression was visualized by *in situ* hybridization and GFP and BrdU signals were visualized by immunofluorescence. The *Shh*, BrdU, and nucleus (Hoechst) visualizations were on the identical section. (A–J) The control vector, pCS-AcGFP, was electroporated into the blastema mesenchyme. The area surrounded by the arrowheads in C shows the area with GFP signals. (F–J) Reconstituted 3D image by Amira. (F1–J1) The proximal view images of the indicated area in F2-J2. The nine sections were extracted from the blastema. *Shh* = orange. *Fgf8* = light green, GFP = yellow, BrdU = light blue, cell density = heatmap. (K–T) The vector, pCS-Shh-p2a-AcGFP, was electroporated into the blastema mesenchyme. The area surrounded by the arrowheads in K and M indicates the area with Shh-p2A-AcGFP signal. (P–T) Reconstituted 3D image by Amira. (P1–T1) The proximal view images of the indicated area in P2-T2. The nine sections were extracted from the blastema. *Shh* = orange. *Fgf8* = light green, GFP = yellow, BrdU = light blue, cell density = heatmap. Scale bars in A and K = 400 µm, in F1 and P1 = approximately 400 µm.

### Digit formation based on the relationship between Shh and Fgf8

Lastly, we investigated the effects of increased cell proliferative activity and cell density on morphogenesis at the sites flanked by *Shh*^+^-*Fgf8*^+^ domains. We hypothesized that the increase in cell density leads to cartilage differentiation since it was shown that higher cell density leads to cartilage differentiation in chicken limb bud culture ^21^. First, we investigated gene expression patterns in the regenerating blastema on both longitudinal and transverse sections, in which the first digit started differentiating (Fig. 9). *Shh* and *Ptc1* were used to visualize SHH distribution (Fig. 9A, B, F). *HoxA13* was used to show the presumptive autopod area (Fig. 9D, I), and the forming cartilage element was shown by *Col2a* (Fig. 9C, H). *Fgf8* is visualized only on the transverse sections due to its characteristic dorsoventrally separated expression pattern (Fig. 9G). The longitudinal sections are adjacent sections, not identical sections. However, based on the boundary between the blastema mesenchymal cells and the epithelium, an approximate merged image was created (Fig. 9E). The merged image shows that the *Col2a*^+^ area appeared near the *Shh^+^* domain within the *HoxA13^+^* domain (Fig. 9E). The transverse sections also confirm the result that the *Col2a^+^* region appeared between the *Shh*^+^ and *Fgf8*^+^ domain within the *HoxA13^+^* area (Fig. 9F–I). This suggests that the first digit is formed in the area between the *Shh*^+^ and *Fgf8*^+^ domain.

**Fig. 9.**
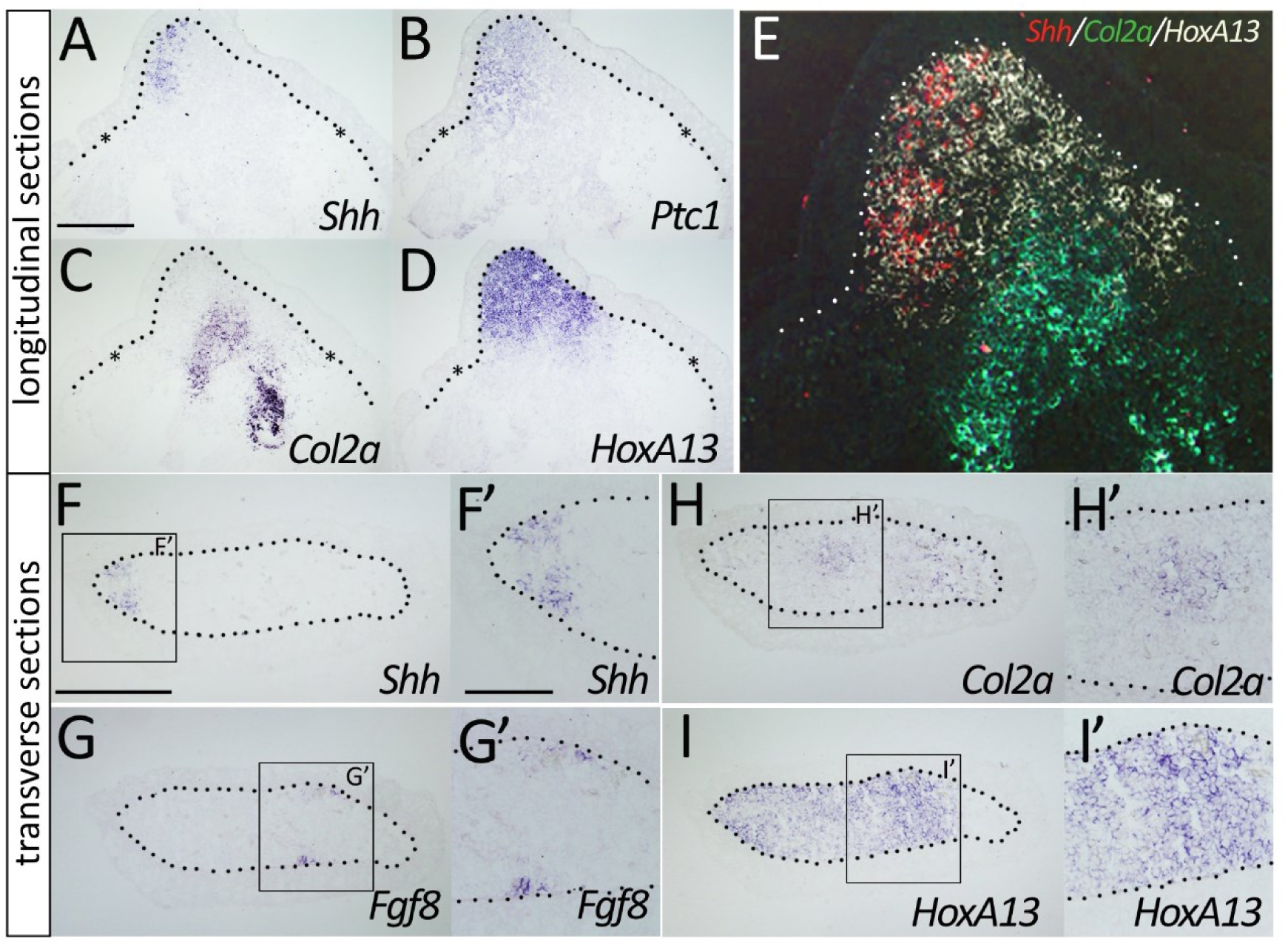
First digit formation and *Shh* and *Fgf8* expression in the blastema at the first digit forming stage. (A–E) *Shh* (A), *Ptc1* (B), *Col2a* (C), and *HoxA13* (D) are visualized on longitudinal sections by *in situ* hybridization. The asterisks indicate the punctuated derma collagen layer. (E) Approximate merged illustration reconstituted from three adjacent sections. The locations of *Shh^+^* cells (red) and *Col2a^+^* cells (green) are projected onto the *HoxA13* (white) section. Scale bars in A = 500 µm. (F–I) *Shh* (F), *Fgf8* (G), *Col2a* (H), and *HoxA13* (I) are also visualized on the transverse sections. F’–I’ are the higher magnification views of F–I. Scale bar in F = 500 µm. Scale bar in F’ = 200 µm. The dotted lines indicate the border of the blastema epithelium. The Posterior is to the left (A–I).

The morphogenetic impact of the variable *Shh*^+^-*Fgf8*^+^ domain in a blastema was assessed in finger morphogenesis. As demonstrated above, the position of the *Shh*^+^-*Fgf8*^+^ domain varies especially in larger blastemas. We also showed that the first digital cartilage appears between the *Shh*^+^ and *Fgf8*^+^ domains. We were curious how the variable *Shh*^+^-*Fgf8*^+^ domain affects digit morphogenesis after the initial digit chondrogenesis occurs. First, the gene expression pattern was revealed after the formation of the first- and second-forming digit cartilages (Fig. 10A-E). We visualized *Shh* expression after first- (digit II) and second-digit (digit I) formation by *in situ* hybridization. Previous reports demonstrated that *Shh* expression was maintained even after digit I and II formation in axolotl limb development^8, 9^. Consistently, *Shh* expression was confirmed in the blastema after digit I and II formation (Fig. 10A, A’, C, C’ E, n = 4/4). *Fgf8* expression was also investigated and found on the adjacent section (Fig. 10B, B’, E). *Fgf8* expression on the ventral side usually disappeared at this stage (n = 4/4). The *Col2a^+^* domain became larger compared to that in Fig. 9H (Fig. 10C, C’) due to the growth of cartilage as the regeneration stage progressed. We also confirmed that the digit I formation, which has been considered to be free from Shh-signaling, occurred in the anterior region (Fig. 10C, C’). *HoxA13* expression indicates that expressions of *Shh*, *Fgf8*, and *Col2a* are in an autopodial region (Fig. 10A-D). To focus on the positional relationship of *Shh*, *Fgf8*, and *Col2a* expression, the spatial positions of *Shh*^+^, *Fgf8*^+^, and *Col2a*^+^ cells were plotted, and merged from the adjacent section, and projected to the section showing *Col2a* expression (Fig. 10E). Considering the relative position of *Shh*^+^, *Fgf8*^+^, and *Col2a*^+^ cells, the location of the *Shh*^+^ and *Fgf8*^+^ domain can be thought to be moved posteriorly as compared to those in the early stage (Fig. 9F–H vs Fig. 10E). This suggests that the aPZ flanked by the *Shh*^+^ and *Fgf8*^+^ domain is gradually shifted posteriorly as digit morphogenesis progresses.

**Fig. 10.**
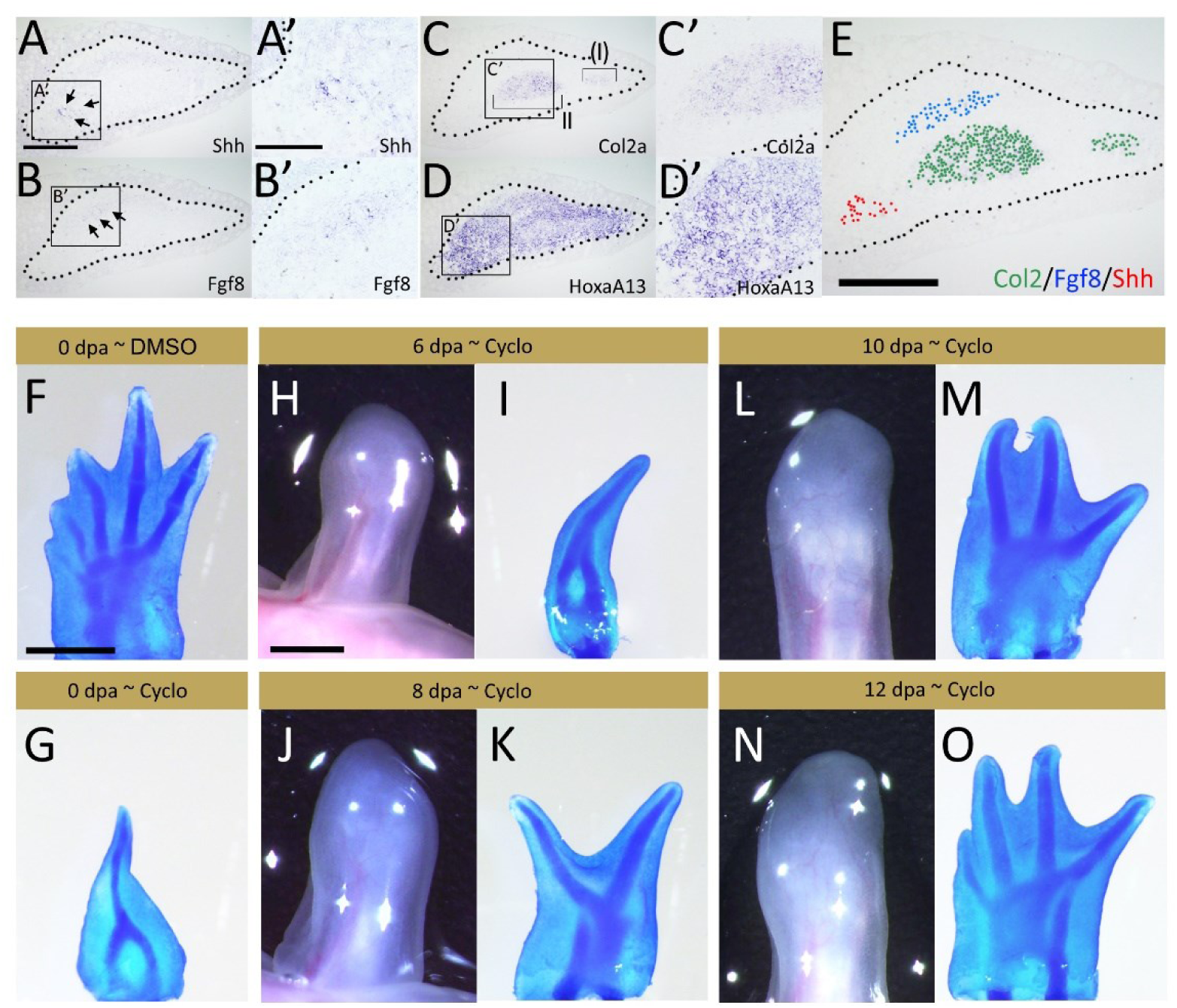
Posterior digit formation and *Shh* and *Fgf8* expression in the blastema at the two-digit stage. (A–E) Gene expression pattern was investigated in the blastema prior to third digit formation. *Shh* (A, A’), *Fgf8* (B, B’), *Col2a* (C, C’), and *HoxA13* (D, D’) were visualized by *in situ* hybridization. (E) Approximate merged illustration reconstituted from three adjacent sections. The locations of *Shh^+^* cells (red) and *Fgf8^+^* cells (blue) are projected onto the *Col2a* (green) section. Scale bars = 500 µm [A, E] and 250 µm [A’]. The dotted lines indicate the border of the blastema epithelium. (F–O) Limb phenotypes produced by cyclopamine exposure with different treatment lengths. (F) Negative control (DMSO treatment). (G) The representative skeletal pattern, which had been treated from the beginning. (H, J, L, N) Blastema morphologies at the start of cyclopamine exposure. (I, K, M, O) Limb morphologies at the end of cyclopamine exposure. Scale bars = 1 mm [F, H]. The Posterior is to the left (A–O).

Lastly, we investigated whether shifting the aPZ toward the posterior side has an impact on posterior digit morphogenesis. We cut off the Shh-signaling at various timepoints after limb amputation and observed the digit formation (Table 1, Fig. 10F–O). Shh-signaling was blocked by cyclopamine treatment. Cyclopamine treatment immediately after limb amputation resulted in spike formation as previously reported whereas DMSO treatment did not induce any defects ^22^ (Fig. 10F, G). However, when the timing of the cyclopamine treatment was delayed, the digit loss number decreased accordingly (Table 1, Fig. 10H–O). A similar observation can be found in axolotl limb development ^23^. This suggests that the posteriorly shifting *Shh*^+^ domain contributes to posterior digit morphogenesis. We further investigated whether exogenous *Shh* and *Fgf8* expression could induce an additional aPZ, leading to additional digit formation (Fig. 11). The experimental scheme is as indicated in Fig. 11A. We electroporated *Shh* and *Fgf8* into the posterior region of the intact limb posterior-most digit. To promote digit morphogenesis, we created a small skin wound in the area, where the introduced gene expression could be confirmed. The introduced gene expression could be expected as a temporal expression. Thus, we performed additional electroporation 13 days post first-electroporation (dpe) (Fig. 11A). The electroporation successfully delivered the exogenous gene expression locally (Fig. 11B). The time course after the electroporation was shown in Fig. 11C. When *Gfp* expression was delivered in the posterior region, no bud formation could be confirmed and no extra-digit formation could be confirmed except for one case (Fig. 11C, left, Table 2, Fig. 11D, n = 23/24). When *Fgf8* was introduced, most limbs grew a bud, but the formed bud did not maintain its growth (Fig. 11E, Table 2, n = 7/14). However, we could observe digit formation in a few cases (Table 2, n = 3/14). For *Shh* electroporation, most cases had no reaction (Fig. 11F, n = 13/16), and a few cases showed bud formation or extra-digit formation (Table 2, n=3/16). When both *Shh* and *Fgf8* were co-electroporated, the induction rate was largely changed. In contrast, when *Shh* and *Fgf8* were electroporated, bud was usually observable by 13 dpe (Fig. 11C, right). Most of the buds induced by the exogenous *Shh* and *Fgf8* reached extra-digit formation (Fig. 11G, n = 9/15). We also found that the additional *Fgf2* expression led to much more effective-extra digit induction. As *Shh*, *Fgf8*, and *Fgf2* were introduced in the posterior region of the autopod, 1 to 4 digit(s) were exogenously induced (Fig. 1H, I, Table 2, n = 13/13). Given the mitogenic role of FGF2, additional *Fgf2* expression might contribute to creating more space for digit formation. Lastly, we note that the extra-digit induction could be induced not only in the posterior-most region, but also in any places within the autopodial region (Sup. Fig. 3). The exogenous activities of Shh- and Fgf-signaling in the anterior or inter-digital region gave rise to the extra digit formation (Sup. Fig. 3). These findings suggest that the exogenously introduced *Shh* and *Fgf8* expression cause extra-aPZ formation, leading to digit formation.

**Fig. 11.**
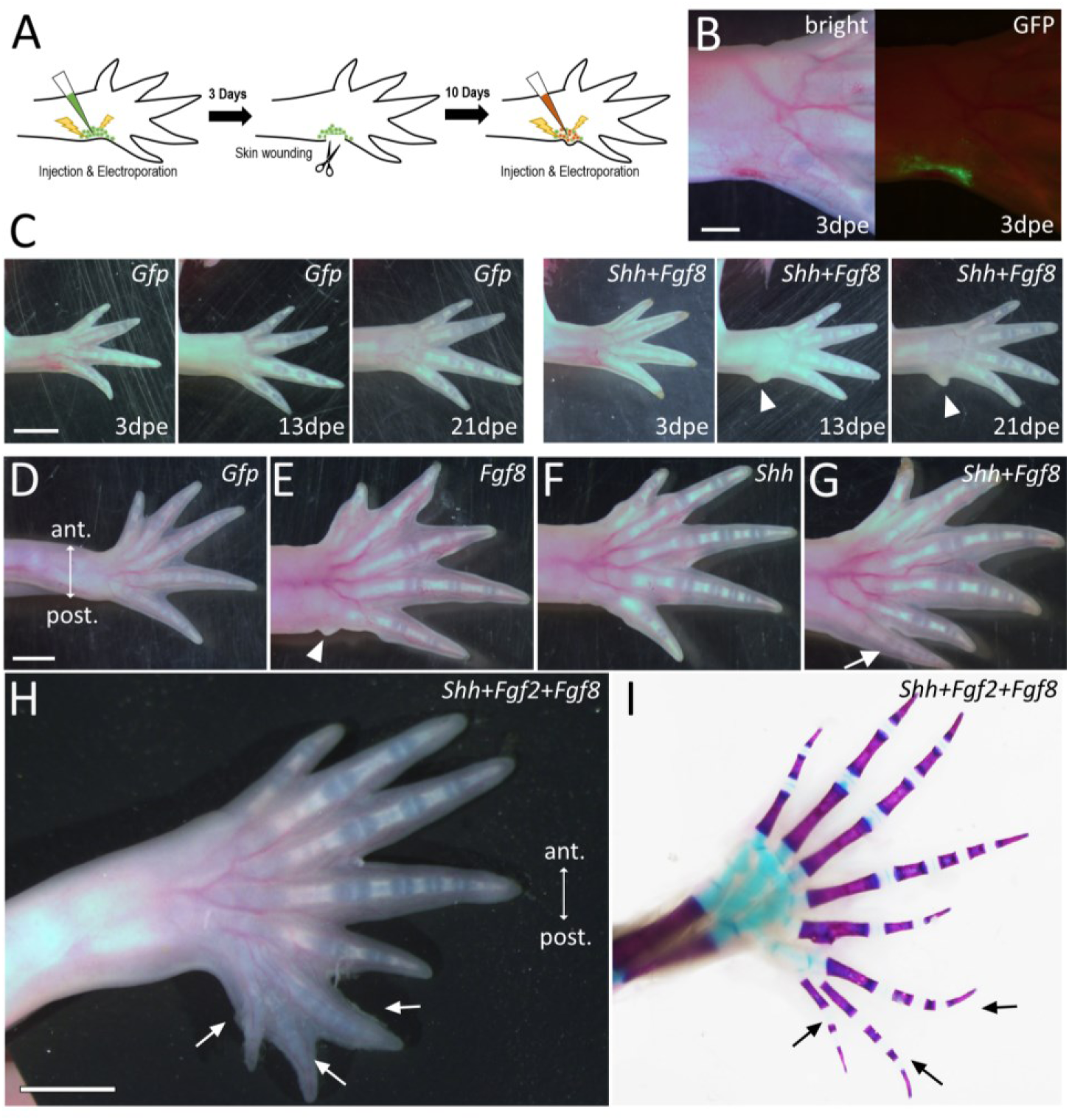
Additional digit formation caused by exogenous Fgf and Shh signaling in the posterior hand/leg. (A) Experimental scheme (hind limb). (B) The dorsal view of the electroporated hand 3 days after the electroporation (before skin wounding). The GFP signals could be found restrictedly in the posterior end of the limb. (C) The time-course of additional digit formation (forelimbs). The left side shows CTRL (pCS-AcGFP electroporation) and the right side shows the condition under the *Shh* and *Fgf8* electroporation. (D–H) The typical digit phenotypes of the electroporated limbs (hind limbs). (D) The control limb (pCS-AcGFP electroporated). In most cases, no extra digit was induced. (E) The *Fgf8* electroporated limb. Only a bud was induced in most limbs (arrowhead). (F) The *Shh* electroporated limb. In most cases, no extra digit or bud was induced. (G) The *Shh* and *Fgf8* co-electroporated limb. Additional digit formation was induced in most of the limbs (arrow). (H) The *Shh*, *Fgf2*, and *Fgf8* co-electroporated limb. Multiple additional digit formation was induced in the posterior region (arrows, hind limb). (I) The skeleton pattern of the sample is shown in H. Scale bar in B, D, and H = 2mm.

**Table 1.** Digit loss in cyclopamine-treated blastemas with different application time.

| Number of lost digit | 0 | 1 | 2 | 3 | 4 |
| --- | --- | --- | --- | --- | --- |
| 0 dpa- cyclo. FL. |  |  |  | 6 |  |
| 0 dpa- cyclo. HL. |  |  |  |  | 6 |
| 6 dpa-cyclo. FL. |  |  |  | 6 |  |
| 6 dpa-cyclo. HL. |  |  |  | 3 | 3 |
| 8 dpa-cyclo. FL. |  |  | 6 |  |  |
| 8 dpa-cyclo. HL. |  |  |  | 6 |  |
| 10 dpa-cyclo. FL. | 4 | 2 |  |  |  |
| 10 dpa-cyclo. HL. |  |  | 6 |  |  |
| 12 dpa-cyclo. FL. | 6 |  |  |  |  |
| 12 dpa-cyclo. HL. | 1 | 3 | 2 |  |  |
| 0 dpa-DMSO FL. | 6 |  |  |  |  |
| 0 dpa-DMSO HL. | 6 |  |  |  |  |

**Table 2.** The number of additional digit formation caused by intact limb electroporation.

|  | +0 | bud only | +1 | +2 | +3~4 |
| --- | --- | --- | --- | --- | --- |
| *Shh+Fgf2+Fgf8* (N=13) | 0<sub>/13</sub> | 0<sub>/13</sub> | 2<sub>/13</sub> | 5<sub>/13</sub> | 6<sub>/13</sub> |
| *Shh + Fgf8* (N=15) | 4<sub>/15</sub> | 2<sub>/15</sub> | 6<sub>/15</sub> | 3<sub>/15</sub> | 0<sub>/15</sub> |
| *Fgf8* (N=14) | 4<sub>/14</sub> | 7<sub>/14</sub> | 3<sub>/14</sub> | 0<sub>/14</sub> | 0<sub>/14</sub> |
| *Shh* (N=16) | 13<sub>/16</sub> | 2<sub>/16</sub> | 1<sub>/16</sub> | 0<sub>/16</sub> | 0<sub>/16</sub> |
| *Gfp* (N=24) | 23<sub>/24</sub> | 0<sub>/24</sub> | 1<sub>/24</sub> | 0<sub>/24</sub> | 0<sub>/24</sub> |

## Discussion

### Characteristic expression pattern of Shh and Fgf8 in regenerating limb blastemas guarantees consistent morphogenesis over the size differences

The present study shows that regenerating axolotl limbs have a characteristic *Shh* and *Fgf8* expression pattern during limb regeneration. As in tetrapods, *Shh* is expressed in the posterior margin of a developing axolotl limb bud ^9^. However, the *Shh* expression is not necessary to be restricted in the posterior margin in the larger size of a blastema (Fig. 2E, F). This segregated *Shh* expression pattern from the blastema epithelium is quite characteristic since *Shh* expression in a developing limb bud is maintained by epidermal signals in other species^7, 8, 9^. *Shh* is maintained by FGFs secreted from the AER in amniotes ^11^. The *Shh*^+^ domain restriction to the posterior margin is controlled by *dHand* (*Hand2*), *Tbx2*, *Gremlin*, and *Alx4* ^24, 25, 26^. In axolotl limbs, the expression pattern of *Tbx2*, *dHand*, *Alx4*, and *Gremlin* was already reported^7, 8, 27^. Among those, no characteristic expression pattern appears to be found as compared to other vertebrates. Only *Fgf8* shows an apparent difference among the genes reported to play roles in *Shh* spatial restriction. *Fgf8* is expressed not in the epidermis but in the anterior mesenchyme of a blastema ^7^ (Fig. 2C, D, G, H). It was reported that the mesenchymal *Shh* and *Fgf8* have been demonstrated to have a positive feedback loop via *Gremlin* ^7^. The present study also clarified the positive interaction between them (Sup. Fig. 2A). This mesenchymal *Fgf8* expression and the positive interaction of *Shh* and *Fgf8* may account for the unique *Shh* expression domain. As in amniotes, *Shh* is likely maintained by *Fgf8* in axolotl limbs. However, unlike in amniotes, the *Fgf8*^+^ domain is in the anterior side of the mesenchyme. The position of the *Fgf8*^+^ domain is varied in blastemas, and a larger size provides more space to be positioned from the center to the anterior side of a blastema (Fig. 4D). Given the mutual dependence between *Shh* and *Fgf8* expression and constant SHH diffusion in the different sizes of blastemas (Fig. 5H–K), it is reasonable that the anterior shift of the *Shh*^+^ domain leads to an anterior shift of the *Fgf8*^+^ domain (Fig. 7Q–X, Fig. 12A). Similarly, as the *Shh*^+^ domain takes a posterior shifted position, the *Fgf8*^+^ domain takes a posterior position accordingly (Fig. 10A, B, Fig. 12A). Furthermore, the constant working range of SHH regardless of the blastema size likely allows SHH reaching to the very anterior end in a quite small blastema. So, the expression region of *Fgf8* can be expanded to the anterior end if a blastema size is small enough. We also demonstrated that the active cell proliferation was formed in the area flanked by *Shh*^+^ and *Fgf8*^+^ domains. We determine the area as aPZ. The variable *Shh*^+^ and *Fgf8*^+^ domain make the positioning of the aPZ variable as well. The higher mitotic activity in the aPZ might be supported by the mitogenic activity of SHH and FGF8, respectively, as well as the coordinated higher cell proliferation activity of both (Sup. Fig. 2B). This characteristic *Shh* and *Fgf8* regulation and associated aPZ formation can account for the ability of limb regeneration in blastemas with varied sizes. In other words, the variable *Shh^+^* and *Fgf8^+^* domain and associated aPZ formation in axolotl limb regeneration might guarantee consistent morphogenesis in the limb regeneration induced in different sizes of animals.

**Fig. 12.**
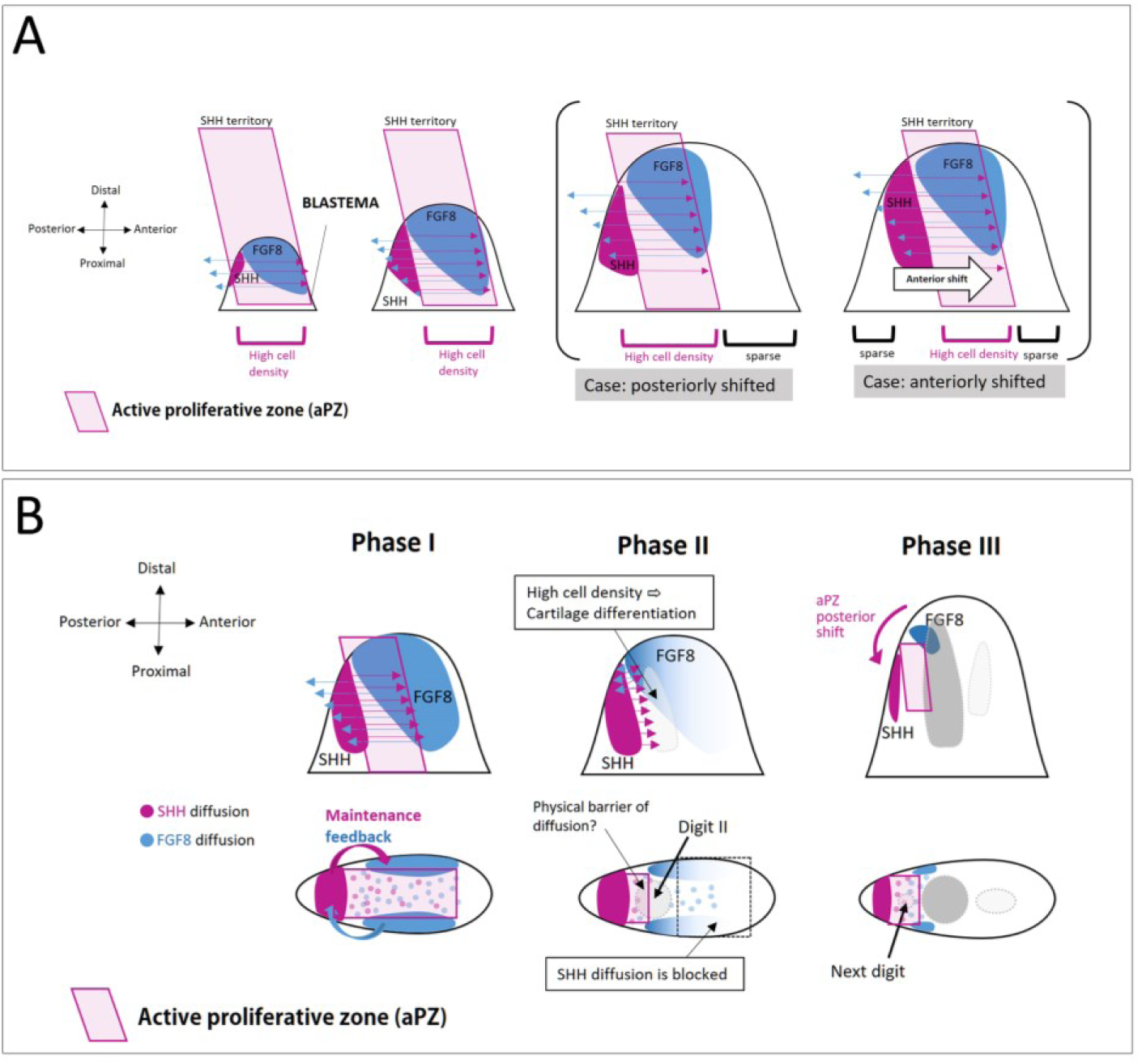
Schematic diagram of the proposed limb patterning mechanism in axolotl limb regeneration. (A) SHH and FGF8, which are interdependent, are within their respective diffusion ranges in a small blastema. As a blastema becomes bigger, regions outside the diffusion range of SHH are produced. As the diffusion range of SHH is constant due to physiological characteristics, it is not affected by the size of the blastema. In a large blastema, the region of SHH-FGF8 is variable. This inconsistent positioning arises from the inconsistent contribution of cells from anterior and posterior stump tissues to a blastema. The area under the influence of SHH and FGF8 shows higher mitogenic activity in a blastema. (B) A model of the anterior-posterior digit formation in an axolotl limb. Owing to the higher mitogenic activity in the area where they can receive both SHH and FGF8, cartilage differentiation is triggered in accordance with increasing cell density. In contrast, FGF8 also shows mitogenic activity. Thus, cartilage differentiation occurs slightly later in the anterior side flanked by FGF8. Differentiating cartilage should be rich in the extracellular matrix and be a physical barrier to SHH and FGF8. Then, FGF8 is maintained in the smaller area, where SHH can be reached, resulting in the posterior shift of the SHH-FGF8 domain. The next presumptive mitogenic point is formed between the shifted SHH and FGF8 domains. This sequence might explain urodele-specific anterior–posterior-ordered digit formation.

Research on the relationship between limb regeneration and size in axolotls has only recently begun to be focused on, and insights into limb size and regeneration have been reported ^28^. It was shown that the amount of nerves in a blastema controls the size of the regenerated limb. In a larger limb, more nerves are present at the amputation surface, and the presence of more nerves contributes to the regeneration of a larger limb, while smaller limbs have a relatively smaller amount of nerves and can only regenerate smaller limbs. This is consistent with experimental findings in axolotls, in which repeated excision of regenerative buds results in progressively smaller regenerating bodies, which have a reduced amount of nerves ^29^. These two studies suggest the significant effects of nerves on size regulation in axolotl limb regeneration. How the nerves regulate the size control is unknown. Given that the nerves can be the source of FGF8 during axolotl limb regeneration^30, 31^, nerve presence might be involved in aPZ regulation. Although it is known that the nerves project axons to a limb, the regularity of the axon route in a blastema has not yet been reported. It is conceivable that FGF8 released from axons has no clear regularity. This might be related to the relatively poor regularity of the *Fgf8*^+^ domain of the blastema mesenchyme compared to the stability of *Shh*^+^ domains (Fig. 4). As the axon routes are varied, the *Shh* and *Fgf8* expression might be affected, leading to variation of the aPZ. The involvement of nerves in aPZ regulation and size regulation in axolotl limb regeneration should be carefully investigated in the future.

### Cell memory and Shh/Fgf8 expression in axolotl blastema

Blastema cells are derived from fully differentiated tissues. This is one of the biggest differences between a developing limb bud and a regenerating blastema. Mesenchymal cells of a developing limb bud are derived from undifferentiated cell sources, such as the lateral plate mesoderm. In contrast, blastema cells are induced from differentiated tissues, such as dermis ^32^. Cell origin must have a great influence on limb morphogenesis, even though a developing limb bud and a regenerating blastema exhibit similar genetic cascades in morphogenesis. We have previously demonstrated that the gene expression pattern reflects cell origin ^33^. *Shh*-expressing cells only emerge from posteriorly derived cells. Thus, it is very likely that *Fgf8*-expressing cells are derived from anterior tissues. This “positional memory” of cells would be unique to limb regeneration compared to limb development. In limb development, it is considered that recognition of a location of a cell is sequentially induced following the developmental program. In contrast, in limb regeneration, the positional memory is in cells before amputation, and it is activated after amputation. The entities of the positional memory have not yet been clarified. However, this “pre-determined cellular memory” may be important for limb regeneration. In contrast to limb development, the linages of cells that form a blastema, rather than induction by secreted factors, might contribute more to the determination of spatial coordinates within a blastema. The dermis is a major contributor to blastema ^34^. The quantitative contribution from dorsal/ventral/anterior/posterior tissues has not yet been carried out. However, it is very likely that the cell contribution from each orientation is varied. Thus, the ratio of blastema cells derived from anterior/posterior tissues may be uneven in each blastema. This may be the cause of the variation in the position of the *Fgf8* expression domain in the blastemas (Fig. 4D). If the contribution of anterior cells to a blastema is large, the anteriorly derived cells will dominate in a blastema. This might lead to posteriorly shifted *Shh*-*Fgf8* expression. Similarly, if the posterior cell contribution is dominant, an anteriorly-shifted *Shh*-*Fgf8* expression would be expected (Fig. 7Q–X, Fig 12A). The unique limb morphogenesis accorded by variable *Shh*-*Fgf8* expression guarantees robust limb regeneration in such a varied cell contribution. Further research is needed to better understand the memory of the orientation and the contribution rate.

It is likely that *Shh*^+^ cells and *Fgf8*^+^ cells are derived solely from posterior tissues and anterior tissues, respectively, as mentioned above. This might be the reason why the *Shh*^+^ and *Fgf8*^+^ domains were not overlapped. As reported, *Shh* and *Fgf8* are mutually dependent. However, the ability of the re-expression of *Shh* or *Fgf8* is predetermined by the origin of the cell. Our previous report showed that the spatial location of cells before amputation dictates gene expression in a regeneration blastema ^33^. Considering this, the *Shh*^+^ domain would be restricted within the area that, the posterior-derived cells occupy, under the influence of FGF8. The *Fgf8*^+^ domain would be restricted in a similar manner. The relationship between the expression of other genes regulating *Shh* or *Fgf8* and their cellular originis still unclear. Further analysis is needed in this regard.

### Variable aPZ leads to digit formation in a blastema

The present study also provides insights into axolotl-specific digit morphogenesis. SHH and FGF8 have synergic cell proliferative activity, forming the aPZ in a blastema (Sup. Fig. 2B). As mentioned, the aPZ is variable in the position since the *Shh*^+^/*Fgf8*^+^ domain varies in a larger blastema. We also clarified that cell density was increased in the aPZ (Figs. 6 and 7). However, it remains unknown whether the increasing cell density and higher mitogenic activity in the aPZ have a causal relationship or just a correlation. Our results suggest that digit-cartilage differentiation occurs in the aPZ (Fig. 9). In evolutionary morphology, the formation of the first digit occurs near the *Shh*^+^ domain ^35^. From this perspective, the mechanism by which the formation of the first digit is induced in the vicinity of the *Shh*^+^ domain can be considered to be conserved in the axolotl. However, the aPZ was shown to be variable in position especially in a larger size of a blastema (Fig. 4G, Fig. 7I – X). Furthermore, the aPZ is gradually shifted posteriorly due to the mutual maintenance mechanism as digital differentiation progresses (Fig. 10). The mechanism of this posterior shifting of the aPZ is still largely unknown. However, it is very likely that the differentiating cartilage becomes a physical block to SHH. Then, the *Fgf8*^+^ domain is only maintained in the posterior tip since only the posterior tip of the *Fgf8*^+^ domain can receive SHH from the posterior side of a blastema (Fig. 12B). This continuous posterior shifting of the aPZ would contribute to the formation of axolotl digits anterior to posterior order. How to terminate the aPZ shifting after posterior-most digit formation remains unknown. However, if SHH and FGF8 were extendedly and exogenously expressed, extra digits beyond the normal posterior-most digit could be formed (Fig. 11). This suggests that the termination of *Shh* and *Fgf8* expression in normal limb morphogenesis is not absolute and irreversible, rather that it fades leaving digit formation competency in the more posterior region. Digit formation in urodele amphibians is known to differ from that of other tetrapods ^13^. The mechanism of digit formation suggested in this study is quite distinctive, although the principle that the first digit is formed in the vicinity of the *Shh*^+^ domain is conserved. The digit formation mechanism revealed and proposed in this study may explain the characteristic digit formation in urodele amphibians.

In conclusion, we propose a novel axolotl limb morphogenetic mechanism that guarantees consistent limb morphogenesis, regardless of blastema size. The *Shh*^+^ and *Fgf8*^+^ domain is maintained in a mutually dependent manner and gives rise to the aPZ in the axolotl blastemas between them. Due to the consistent diffusion distance of the SHH, regardless of blastema size, the aPZ can be maintained at a relatively consistent size. In a larger blastema, the position of the aPZ within the blastema is relatively variable, but the expected size of the aPZ remains consistent due to the consistent SHH diffusion distance. Regardless of the aPZ positioning in a blastema, the aPZ increases cell proliferation and cell density within a consistent region, leading to the first phalangeal differentiation. Thanks to the variable positioning mechanism of the aPZ, the posterior phalangeal differentiation is given by the posterior-shifting aPZ mechanism. As long as the aPZ is maintained, axolotl digital morphogenesis is maintained. This unique limb morphogenesis mechanism could reasonably account for the unique limb regeneration ability of axolotls throughout their life and digit morphogenesis.

## Materials and Methods

### Animals and Surgery

For most experiments, axolotls (*Ambystoma mexicanum*) with a nose-to-tail length of 8 to 15 cm obtained from private breeders and housed in aerated water at 22°C were used. A larger animal (25cm) was used to show its regeneration process and smaller animals (5 cm) were used for chemical treatment. Before all surgeries, axolotls were anesthetized using MS-222 (Sigma-Aldrich, St. Louis, Missouri) for about 10 minutes (depending on the animal size). Limbs were amputated at the middle of the zeugopod. All animal usage was approved by the Animal Care and Use Committee, Okayama University (#580 to A.S), and all animal experiments were conducted following the guidelines of Okayama University.

### Sectioning and histological staining

Samples were fixed with 4 % paraformaldehyde for 1 day at room temperature and embedded in O.C.T. compound (Sakura Finetek, Tokyo, Japan) following 30% sucrose/PBS treatment for approximately 12 h. Frozen sections with a thickness of 10 μm were prepared using a Leica CM1850 cryostat (Leica Microsystems, Wetzlar, Germany). The sections were dried under an air dryer and kept at −80°C until use. Standard hematoxylin and eosin (HE) staining was used for histology. To visualize cartilage formation, Alcian blue staining was performed before HE staining. In brief, sections were washed in tap water several times to remove the O.C.T. compound. Then, alcian blue (Wako, pH 2.0) solution was dropped on the section and the slide was incubated for 2 min. The sections were washed twice with tap water, and then HE staining was performed. The stained sections were mounted using Softmount (Wako). For skeletal preparation, samples were fixed with 95% ethanol for 1 h. Then, the cartilage was stained with approximately 0.1% Alcian blue solution (80% ethanol/20% acetic acid) for 2 days at 38°C. Then, the samples were placed in approximate 0.1% Alizarin red solution (2%KOH/ 2% paraformaldehyde) overnight at room temperature. Tissue transparency was achieved by stepwise replacement of the glycerol/KOH solution. Finally, the transparency process was completed with 100% glycerol replacement. Each substitution step took 2 to 5 days. The histological observation was confirmed in multiple samples.

### Flow cytometry

Blastema cells were dispersed using collagenase (Wako Pure Chemical Industries, Osaka, Japan). Filtration was then performed using Cell Strainer (FALCON, 45mm mesh). Cells were suspended in 40% Dulbecco’s modified Eagle Medium (gibco), 50% DDW, 10% FCS, 0.01M HEPES, 3µg/mL gentamycin. The cells were then resuspended in PBS. Cells were dissolved in 1 ml of PBS and nuclei were stained with propidium iodide (PI) (Merck Millipore, Burlington, MA, USA) and analyzed using a flow cytometer (Guava PCA). The data were analyzed using FCSalyzer (https://sourceforge.net/projects/fcsalyzer/). The data was obtained by 3 times independent experimental replicates. The merged plot was shown in Fig. 1G.

### Actin staining

After washing samples with TBST (0.15M NaCl, 0.1M Tris and 0.1% Tween20), Alexa Fluor™ 594 Phalloidin (#A12381, Invitrogen, CA, USA), was added at 1:500 dilution and incubated for 20 min at room temperature. Samples were washed with TBST, and nuclei were visualized by Hoechst33342 (#346-07951, Wako-Fuji film).

### *In situ* hybridization, immunofluorescence and nuclear visualizing

RNA probes for *Fgf8*, *Shh, HoxA13*, *Col1a* and *Col2a* were selected as previously described ^[1–4]^. The axolotl *Ptathed 1* (*Ptc1*) gene was newly isolated. The isolated sequences are from 1016-2989 of AMEX60DD201043057.6 in the axolotl OMICS site (https://www.axolotl-omics.org/). *In situ* hybridization was performed as reported (Satoh et al., 2007). Briefly, samples were treated with proteinase K (10µg/ml) at room temperature for 20 min, and riboprobes were hybridized at 62.5°C. Following hybridization, samples were washed with buffer 1 (50% formamide and 5× saline-sodium citrate (SSC)) twice for 20 min and buffer 2 (50% formamide and 2×SSC) three times for 20 min at 62.5°C. After samples were blocked with 0.5% blocking reagent (#11096176001, Roche) /TBST, an Anti-Digoxigenin-AP, Fab fragments (#11093274910, Roche) were added at 1:1000 dilution and incubated for 2 h at room temperature. Staining was done with NBT (#148-01991, Wako-Fuji film) and BCIP (#05643-11, Nakalai Tesque) in NTMT (100 mM NaCl, 0.1 M Tris pH9.5 and 0.1% Tween20). For *in situ* hybridization and BrdU double staining, *in situ* hybridization was performed first, and immunostaining was subsequently performed. Immunofluorescence of the sections was performed as described in previous reports^[5]^. BrdU antigen retrieval was performed with 2 mol/L HCl for 30 min at room temperature. Anti-BrdU (G3G4, 1:300, DSHB, IA, USA) was used for the primary antibody and anti-mouse IgG Alexa Fluor 488 (A11017, 1:500, Invitrogen, CA, USA) was used for the secondary antibody. Nuclei were visualized by Hoechst33342 (#346-07951, Wako-Fuji film) after performing *in situ* hybridization. Images were captured using an Olympus (Tokyo, Japan) BX51 fluorescence microscope.

### Measurement of the signals on sections

Measurement of gene expression domains was performed using software (Cellsense, Olympus). The width of blastema mesenchyme along with the anteroposterior axis and gene expression domain was measured using a measurement tool on Cellsense. To maintain fairness, all measurements were performed under the same exposure time and magnification.

### Simulations for the spatial distribution of SHH

In Figs. 2J-K, we numerically solved the following partial differential equation for diffusion and linear degradation of SHH using Mathematica software (ver. 12.1.1.0).

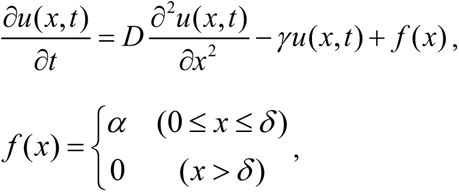

where *u*(*x*, *t*) is the SHH concentration at position *x* and time *t*. *D* and γ are the diffusion constant and degradation rate, respectively, and those values were set so as to 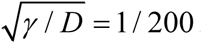. *δ* is the size of SHH expression domain, and we simulated the cases with *δ* = 100 [ μm ] and *δ* = 200 [ μm ]. *α* is the secretion rate of SHH, and we set *α* = 1.0 [a.u.]. A Neumann boundary condition, ∂u / ∂x = 0 , was used at both posterior and anterior ends of a tissue. Regarding tissue size (denoted by *L* ), the cases with *L* = 500 [ μm ] and *L* = 1, 000 [ μm ] were tested. Figures 2J-K show the spatial distribution of SHH at the steady-state.

### SU5402 and cyclopamine treatments

Animals were treated with 10 mL solutions for all chemical treatments. For this experiment, smaller animals were used (5 cm in length). Working stocks of 30 mM SU5402 (#193-16733, Wako) and 8 mM Cyclopamine (#C9710, LKT Labs, Inc.) were made in DMSO (#08904-14, Nakalai Tesque) and diluted to 30 µM and 8 µM in water. Control animals were treated with a solution containing an equal amount of DMSO. The chemical-containing water was refreshed every day (Sup. Fig. 2) or every 2 days (Fig. 10).

### qRT-PCR analysis

RNA preparation for RT-PCR was performed using TriPure Isolation Reagent (Roche). Reverse transcription was performed using PrimeScript™ II Reverse Transcriptase (Takara). The quantitative RT-PCR analysis was performed by StepOne^TM^ (ThermoFisher) and KAPA SYBR FAST qPCR Master Mix (Kapa Biosystems). Primers are described below:

*EF1α* forward, AACATCGTGGTCATCGGCCAT
*EF1α* reverse, GGAGGTGCCAGTGATCATGTT
*Shh* forward, GCTCTGTGAAAGCAGAGAACTCG
*Shh* reverse, CGCTCCGTCTCTATCACGTAGAA
*Fgf8* forward, CGAAGGATGGTACATGGCCT
*Fgf8* reverse, TATCGCGGTACGGAATGTCG
*Pcna* forward, ATGTTTGAGGCTCGCCTGGT
*Pcna* reverse, GCAGCGGTACGTATCGAAGC

### Cell culture and protein addition

Harvested blastemas were removed from the epithelial tissues from blastemas in 1% EDTA/PBS using forceps. Only mesenchyme tissues were used for the cell culture. After washing with PBS, the tissues were broken into smaller pieces. Obtained mesenchyme cells were suspended in culture medium (50% water, 40% GlutaMax-DMEM, 10% FCS and 0.01M HEPES (pH. 7.5), and 300 mg/ml Gentamycin). FGF8 (R&D systems, #423-F8) or/and SHH (R&D systems, #461-SH) proteins were added to the culture medium at a concentration of 0.1 µg/ml. The culture medium was refreshed every day.

### 3D reconstruction and distance map

Two or three series of consecutive blastema sections underwent *in situ* hybridization, BrdU immunofluorescence, and nuclei visualization by Hoechst33342. Images were captured using an Olympus BX51 fluorescence microscope and were reconstructed through volume rendering using Amira Software (version 6.4.0, Thermo Fisher Scientific; http://www.fei.com/software/amira-3d-for-life-sciences/) running on an iMacPro (CPU: 2.3 GHz Intel Xeon W, DRAM: 128GB 2666 MHz DDR4, and graphics: Radeon Pro Vega 64 16368 MB; Apple Japan, Tokyo). Gene expression domains were extracted using the segmentation tool on Amira. For the distance map in Fig. 6 and Sup. Data 1 and 2, nuclei were marked manually on Amira, and nuclei coordinate information was obtained. Nuclei coordinate information was divided into four files due to technical limitations. Distances between nuclei were calculated. The five shortest distances were selected and the average of these five distances was calculated. We found that visualization of the average distance from two to four nuclei was problematic, especially around the blastema margin. In contrast, visualization of the average distance from more than six nuclei looked no different. Thus, we decided to utilize the five nearest nuclei to visualize the cell density. The calculations were performed by the following Python script (https://github.com/satohlab-py/cell-density.git; script name = “uptolist”). These calculations were repeated for every divided four files. The obtained four files were connected manually. The average distances were squared to visualize cell density clearly. Subsequently, the nuclei dots were colored depending on the average distance, and then the dots were reconstructed into the 3D image. These tasks were performed by the following Python script (https://github.com/satohlab-py/cell-density.git; script name = “uptograph”) in the Python programming language (Python Software Foundation, https://www.python.org/). BrdU signals were extracted using the segmentation tool on Amira, and merged signals were isolated using the watershed tool on ImageJ software. BrdU coordinate information was obtained using connected components tools on Amira. Hoechst signals were extracted using the segmentation tool, and a heatmap was created by the DistanceMap tool on Amira.

### Electroporation

The animals were anesthetized as described and the DNA solution was injected directly into the regenerating blastema or under the skin. Then, electroporation was performed on the area where the DNA was injected. The conditions of the electroporation were as follows: 20 V, 20 times, 50 ms pulse, 950 ms interval (NEPA21, Nepa gene). The injected DNA were as follows: pCS-AcGFP, pCS2-Shh-p2A-AcGFP, pCS2-Fgf8 and pCS2-Fgf2-p2A-Fgf8-p2A-mCherry. Shh-p2A-AcGFP, Fgf8, and Fgf2-p2A-Fgf8-p2A-mCherry were created as artificial synthetic genes and subcloned into pCS2 Vector. All plasmids were purified with a Genopure Maxi kit (Roche, Basel, Switzerland) and the final concentration was 2 µg/µl.

## Acknowledgments

We are grateful to R. Iwata and T. Satoh for their laboratory work. We thank K. Nishimura for making a prototype of the Python script. Animals were obtained through Hiroshima University Amphibian Research Funding. This work is supported by a JSPS KAKENHI grant-in-aid for scientific research (B) (#20H03264 to AS).

## Author contributions

S.F.: investigation, data collection, data analysis, methodology, writing–review and editing. S.Y.: supportive investigation, writing–review and editing. R.K.: supportive investigation, animal housing, writing–review and editing. Y.M.: simulation, writing. A.S.: investigation, methodology, conceptualization, supervision, writing–review and editing, project administration, funding acquisition.

## Data and materials availability

All data are available in the main text or the supplementary materials. On reasonable request, derived data supporting the findings of this study are available from the corresponding author (AS).

**Supplemental Movie 1.**

The spatial expression pattern of *Shh* (orange) and *Fgf8* (light green) in the small-size blastema. The image is reconstituted from the sections, in which gene expression was visualized by *in situ* hybridization. Light gray shows the boundary between the mesenchyme and the epithelium. Snapshots from this movie are used in Fig. 3E–H.

**Supplemental Movie 2.**

The spatial expression pattern of *Shh* (orange) and *Fgf8* (light green) in the medium-size blastema. The image is reconstituted from the sections, in which gene expression was visualized by *in situ* hybridization. Light gray shows the boundary between the mesenchyme and the epithelium. Snapshots from this movie are used in Fig. 3K–N.

**Supplemental Movie 3**.

The distribution of nuclei (green) and *Shh*^+^ cell (pink) in the blastema. Snapshots from this movie are used in Fig. 6C and E.

**Supplemental Movie 4.**

The spatial expression pattern of *Shh* (orange) and *Fgf8* (light green) in the small-size blastema. The distribution of BrdU signals (light blue) and the cell density heatmap (deep brown to deep gray) is also visualized. Snapshots from this movie are used in Fig. 7E–H.

**Supplemental Movie 5.**

The spatial expression pattern of *Shh* (orange) and *Fgf8* (light green) in the medium-size blastema. The distribution of BrdU signals (light blue) and the cell density heatmap (deep brown to deep gray) is also visualized. Snapshots from this movie are used in Fig. 7M–P.

**Supplemental Movie 6.**

The spatial expression pattern of *Shh* (orange) and *Fgf8* (light green) in the anteriorly shifted medium-size blastema. The distribution of BrdU signals (light blue) and the cell density heatmap (deep brown to deep gray) is also visualized. Snapshots from this movie are used in Fig. 7U–X.

**Supplemental Data 1.**

The cell density heatmap.

**Supplemental Data 2.**

The cell density heatmap. To avoid a clouded image, the proximal three sections were extracted and visualized.

---

**Supplemental Fig. 1.**
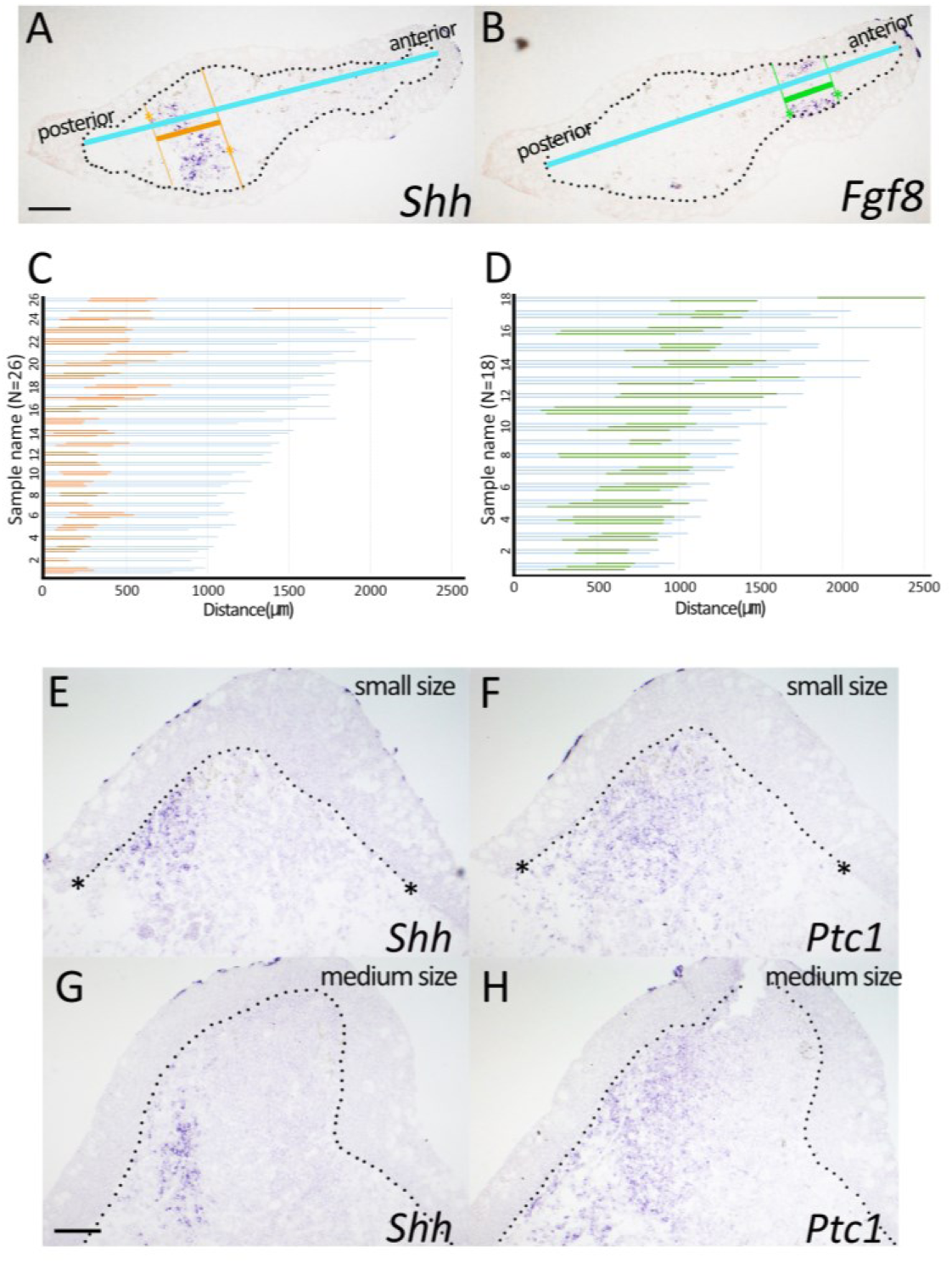
The measurement of the gene expression domain. **(A and B)** The specific measurement method of *Shh* (A) and *Fgf8* (B). The light-blue lines indicate the length of the blastema mesenchyme (anteroposterior direction). The expression domain was indicated by orange (for *Shh*) and green (for *Fgf8*). The dotted lines indicate the border of the blastema epithelium. **(C and D)** The raw data of the *Shh^+^* (C) and *Fgf8^+^* (D) domain widths. The measurement was performed on 2-3 sections derived from an identical blastema. The lines indicate the results from all the measurements of 2-3 sections. The lines were bundled every blastema and displayed. (n=26 (*Shh*), n=18 (*Fgf8*)) (E-H) *Shh* and *Ptc1* expression were visualized in the longitudinal sections of the small (E, F) and medium (G, H) size blastemas. The dotted lines indicate the border of the blastema epithelium. Asterisks indicate the punctuated dermal collagen, indicating the amputation plane. The scale bar in A and G = 200 µm. The Posterior is to the left.

**Supplemental Fig. 2.**
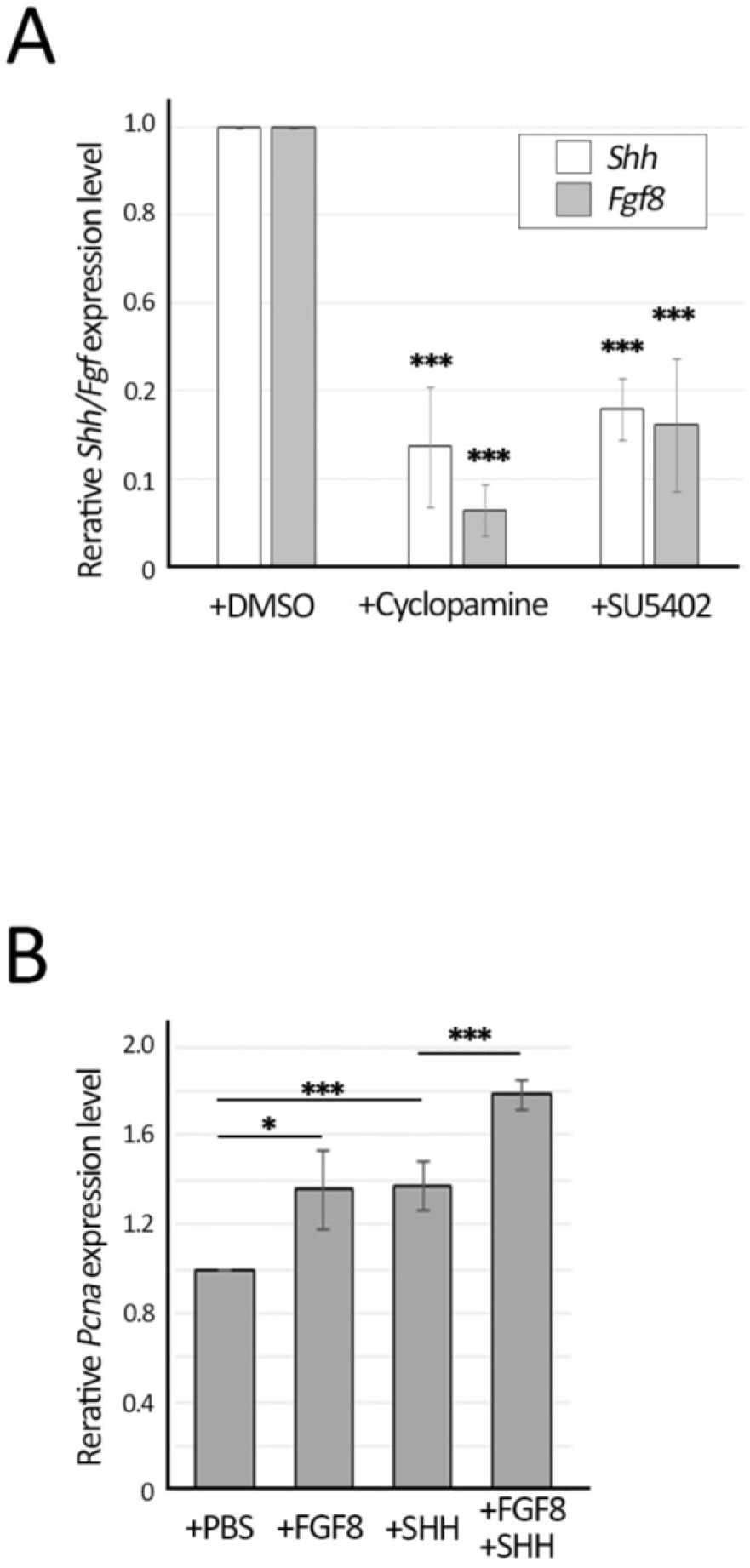
(A) The relative expression level of *Shh* and *Fgf8* was measured by quantitative reverse transcription-polymerase chain reaction (RT-PCR). The statistical analysis was performed using a two-tailed Student’s *t*-test between the control and the samples. *p* values are represented as follows: ***< 0.01. (B) The relative *Pcna* expression level was measured by quantitative RT-PCR. The statistical analysis was performed using a one-way analysis of variance with Ryan’s *post hoc* test. *p* values are represented as follows: ***< 0.01. *<0.05.

**Supplemental Fig. 3.**
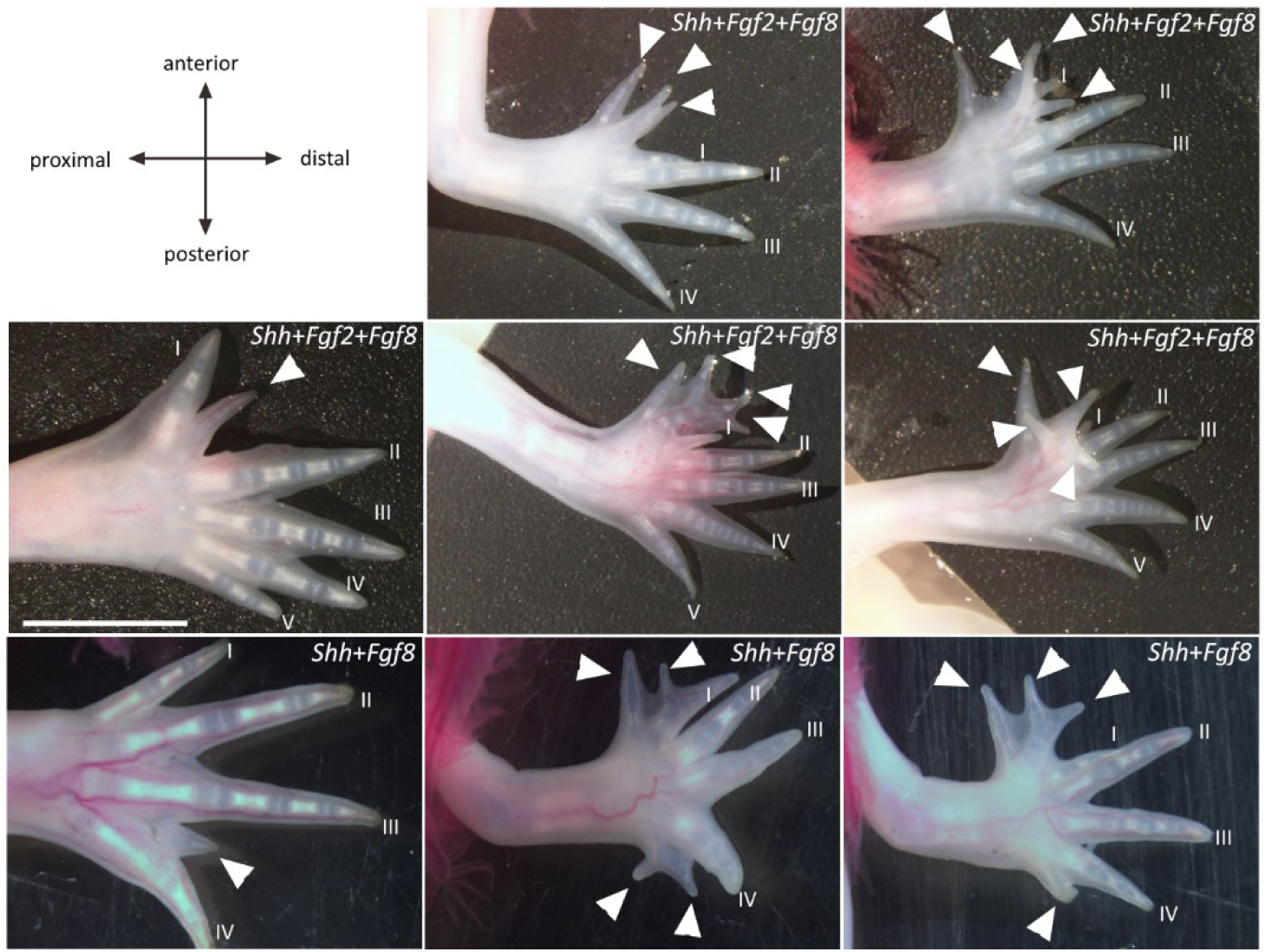
Additional digits induced in the anterior or inter-digital region. Arrowheads indicate the induced extra digits. Roman numbers indicate the digit identity of the original limb. Scale bar = 5mm.

